# Inositol pyrophosphates promote the interaction of SPX domains with the coiled-coil motif of PHR transcription factors to regulate plant phosphate homeostasis

**DOI:** 10.1101/2019.12.13.875393

**Authors:** Martina K. Ried, Rebekka Wild, Jinsheng Zhu, Larissa Broger, Robert K. Harmel, Ludwig A. Hothorn, Dorothea Fiedler, Michael Hothorn

**Author notes:** contributed equally (Martina K. Ried, Rebekka Wild). Leibniz Institute of Plant Biochemistry, 06120 Halle, Germany (Martina K. Ried). Institut de Biologie Structurale (IBS), 38044 Grenoble, France. retired (Ludwig A. Hothorn).

## Abstract

Phosphorus is an essential nutrient taken up by organisms in the form of inorganic phosphate (Pi). Eukaryotes have evolved sophisticated Pi sensing and signalling cascades, enabling them to maintain cellular Pi concentrations. Pi homeostasis is regulated by inositol pyrophosphate signalling molecules (PP-InsPs), which are sensed by SPX-domain containing proteins. In plants, PP-InsP bound SPX receptors inactivate Myb coiled-coil (MYB-CC) Pi starvation response transcription factors (PHRs) by an unknown mechanism. Here we report that a InsP_8_ – SPX complex targets the plant-unique CC domain of PHRs. Crystal structures of the CC domain reveal an unusual four-stranded anti-parallel arrangement. Interface mutations in the CC domain yield monomeric PHR1, which is no longer able to bind DNA with high affinity. Mutation of conserved basic residues located at the surface of the CC domain disrupt interaction with the SPX receptor *in vitro* and *in planta*, resulting in constitutive Pi starvation responses. Together, our findings suggest that InsP_8_ regulates plant Pi homeostasis by controlling the oligomeric state and hence the promoter binding capability of PHRs via their SPX receptors. (173 words)

## Introduction

Phosphorus is an essential building block for many cellular components such as nucleic acids and membranes. It is essential for energy transfer and storage and can act as a signalling molecule. Pro- and eukaryotes have evolved intricate systems to acquire phosphorus in the form of inorganic phosphate (Pi), to maintain cytosolic Pi concentrations, and to transport and store Pi as needed. In green algae and plants, transcription factors have been previously identified as master regulators of Pi homeostasis and Pi starvation responses (PSR)^1, 2^. Phosphorus starvation response 1 (CrPsr1) from Chlamydomonas and PHOSPHATE STARVATION RESPONSE 1 (AtPHR1) from Arabidopsis were founding members of plant unique MYB-type coiled-coil (MYB-CC) transcription factors^3^. PHR transcription factors were subsequently characterized as regulators of Pi starvation responses in diverse plant species^4–6^. In Arabidopsis there are 15 MYB-CCs with PHR1 and PHL1 controlling the majority of the transcriptional Pi starvation responses^7, 8^. Knock-out mutations in *Arabidopsis thaliana PHR1* result in impaired responsiveness of Pi starvation induced (PSI) genes, and perturbed anthocyanin accumulation, carbohydrate metabolism and lipid composition^2, 9,10^. Overexpression of *AtPHR1* causes elevated cellular Pi concentrations and impacts the transcript levels of At*PHO2,* which codes for an E2 ubiquitin conjugase involved in PSR, via increased production of its micro RNA miR399d^9,11^. PHR binds to a GNATATNC motif (P1BS), found highly enriched in the promoters of PSI genes and in other *cis*-regulatory motifs, activating gene expression^2,12^. AtPHR1 is not only implicated in Pi homeostasis, but also in sulphate, iron and zinc homeostasis as well as in the adaption to high-light stress^13–16.^ Moreover, AtPHR1 shapes the plant root microbiome by negatively regulating plant immunity^17^.

AtPHR1 and OsPHR2 have been previously reported physically interact with stand-alone SPX proteins^18–21^, additional components of PSR in plants^21–24^. SPX proteins may regulate PHR function by binding to PHRs under Pi sufficient condition, keeping the transcription factor from entering the nucleus^25–27^. Alternatively, binding of SPX proteins to PHRs may reduce the ability of the transcription factors to interact with their promoter core sequences^19, 20, 25, 26, 28^. Two mechanisms were put forward regarding the regulation of the SPX – PHR interaction in response to changes in nutrient availability: SPX domains were proposed to act as direct Pi sensors, with the SPX – PHR interaction occurring in the presence of millimolar concentrations of Pi^19, 20^. Alternatively, the integrity of the SPX – PHR complex could be regulated by protein degradation. Indeed, SPX degradation via the 26S proteasome is increased under Pi starvation^25, 26, 29^.

Fungal, plant and human SPX domains^30^ have been independently characterized as cellular receptors for inositol pyrophosphates (PP-InsPs), which bind SPX domains with high affinity and selectivity^31, 32^. PP-InsPs consist of a fully phosphorylated *myo*-inositol ring, carrying one or two pyrophosphate groups at the C1 and/or C5 position, respectively^33^. In plants, inositol 1,3,4-trisphosphate 5/6-kinase (ITPK) catalyzes the phosphorylation of phytic acid (InsP_6_) to 5PP-InsP_5_ (InsP_7_ hereafter)^34^. The diphosphoinositol pentakisphosphate kinases VIH1 and VIH2 then generate 1,5(PP)_2_-InsP_4_ (InsP_8_ hereafter) from InsP ^32, 35–37^. Plant diphosphoinositol pentakisphosphate kinases have been genetically characterized to play a role in jasmonate perception and plant defence responses^36^ and, importantly, in nutrient sensing in Chlamydomonas^38^ and Arabidopsis^32, 37^. *vih1 vih2* double mutants lack the PP-InsP messenger InsP_8_, over accumulate Pi and show constitutive PSI gene expression^32, 37^. A *vih1 vih2 phr1 phl1* quadruple mutant rescues the *vih1 vih2* seedling phenotypes and displays wild type like Pi levels, suggesting that VIH1, VIH2, PP-InsPs and PHRs are part of a common signalling pathway^37^. In line with, the AtSPX1 – AtPHR1 interaction is reduced in *vih1 vih2* mutant plants when compared to wild type^32^. Thus, biochemical and genetic evidence implicates InsP_8_ in the formation of a SPX – PHR complex^32, 37^. Cellular InsP_8_ pools are regulated by nutrient availability at the level of the VIH enzymes themselves. Plant VIH1 and VIH2 and diphosphoinositol pentakisphosphate kinases from other organisms are bifunctional enzymes, with an N-terminal kinase domain that generates InsP_8_ from InsP_7_, and a C-terminal phosphatase domain that hydrolyses InsP_8_ to InsP_7_ and InsP ^37, 39, 40^. The relative enzymatic activities of the two domains are regulated in the context of the full-length enzyme: Under Pi starvation, cellular ATP levels are reduced, leading to a reduction of the VIH kinase activity, and a reduction of InsP ^32, 37^. Pi itself acts as an allosteric regulator of the phosphatase activity^37, 39^. Thus, under Pi sufficient growth conditions InsP_8_ accumulates and triggers the formation of a SPX – InsP_8_ – PHR complex. Under Pi starvation, InsP_8_ levels drop and the complex dissociates^41^.

How the InsP_8_ bound SPX receptor inactivates PHR function remains to be understood at the mechanistic level. It has been previously reported that AtPHR1 binds P1BS as a dimer^2^. Addition of SPX domains reduces the DNA binding capacity of PHRs as concluded from electophoretic mobility shift assays (EMSA)^19, 20, 25^. Qi and colleagues reported that AtPHR1 recombinantly expressed as a maltose-binding protein (MBP) fusion protein forms monomers in solution and binds DNA. This process that can be inhibited by preincubating the recombinant transcription factor with AtSPX1 in the presence of high concentrations of InsP ^28^. A recent crystal structure of the AtPHR1 MYB domain in complex with a promoter core fragment supports a dimeric binding mode of MYB-CC transcription factors^42^. Here we investigate the oligomeric state of PHRs, their DNA binding kinetics, and the targeting mechanism of the interacting SPX receptors.

## Results

### PP-InsPs trigger AtSPX1 - AtPHR1 complex formation in yeast

The interaction of AtSPX1 with AtPHR1 has been previously characterized in yeast-2-hybrid assays^19^. We reproduced the interaction of full-length AtPHR1 and AtSPX1 (Fig. 1a) and verified that all four stand-alone AtSPX proteins (AtSPX1 – 4) interact with a AtPHR1 fragment (AtPHR1^226 - 360^) that contains the MYB domain as well as the CC domain in yeast (Supplementary Fig. 1a). This is in line with previous findings, reporting interaction of SPX domains with larger PHR fragments also containing the MYB and CC domains (AtSPX1 – AtPHR1^208–362^ and OsSPX1/2 – OsPHR2^231–426^)^19, 20^.

**Fig. 1.**
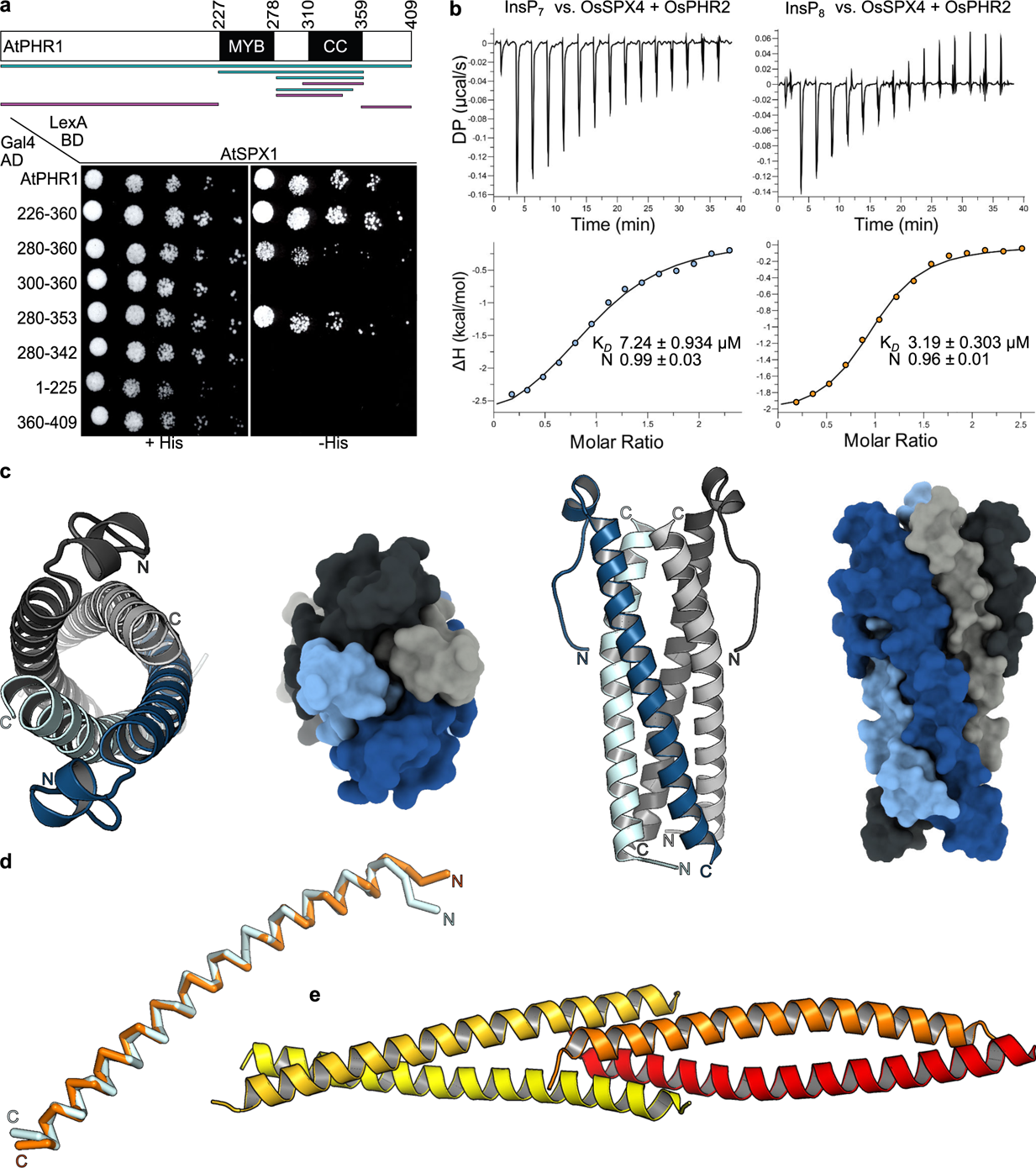
AtSPX1 recognizes the AtPHR1 CC domain that crystallizes as a tetramer. **a** Yeast co-expressing different AtPHR1 deletion constructs fused to the Gal4-activation domain (AD; prey) and full-length wild-type AtSPX1 fused to the LexA-binding domain (BD; bait) were grown on selective SD medium supplemented with histidine (+ His; co-transformation control) or lacking histidine (-His; interaction assay) to map a minimal fragment of AtPHR1 sufficient for interaction with AtSPX1. Shown are serial dilutions from left to right. A schematic overview of the tested interacting (in cyan) and non-interacting (in magenta) AtPHR1 fragments is shown alongside (MYB, DNA binding domain; CC, coiled-coil domain). **b** Isothermal titration calorimetry assays of InsP_7_ (400 µM 5PP-InsP_5_; left panel) and InsP_8_ (500 µM 1,5(PP)_2_-InsP_4_; right panel) binding to OsSPX4 – OsPHR2 (30 µM), respectively. Raw heats per injection are shown in the top panel, the bottom panel represents the integrated heats of each injection, fitted to a one-site binding model (solid line). The insets show the dissociation constant (*K*_d_) and binding stoichiometry (N) (± fitting error). **c** Ribbon and surface diagrams of the AtPHR1 CC four-stranded anti-parallel tetramer. Helices contributing to the dimer interface are shown in light- and dark-blue, respectively. Corresponding, symmetry-related helices completing the tetramer are shown in light and dark-grey. **d** Structural superposition of two core CC helices from AtPHR1 (C_ɑ_ trace, in light blue) and ScCtp1 (PDB-ID 4X01, in orange)^47^. R.m.s.d. is ∼1 Å comparing 45 corresponding C_ɑ_ atoms. **e** Ribbon diagram of the ScCtp1 dimer-of-dimers CC domain, with contributing helices colored from yellow to red.

We next tested if the SPX – PHR interactions observed in yeast are mediated by endogenous PP-InsPs. The putative PP-InsP binding surface in AtSPX1 was mapped by homology modeling, using the *Chaetomium thermophilum* GDE1 – InsP_6_ complex structure (PDB-ID 5IJJ) as template^31^. We replaced putative PP-InsP binding residues from the previously identified <u>P</u>hosphate <u>B</u>inding <u>C</u>luster (PBC: AtSPX1^Y25, K29, K139^) and Lysine (<u>K</u>) <u>S</u>urface <u>C</u>luster (KSC: AtSPX1^K136, K140, K143^)^31^ with alanines (Supplementary Fig. 1b). The resulting AtSPX1^PBC^ and AtSPX1^KSC^ mutant proteins failed to interact with AtPHR1^226-360^ in yeast-2-hybrid assays, while mutation of a conserved lysine residue outside the putative PP-InsP binding site (SC: structural control, AtSPX1^K81^) to alanine had no effect (Supplementary Fig. 1b). We next deleted the known yeast PP-InsP kinases Vip1, which converts InsP_6_ to 1PP-InsP_5_ and 5PP-InsP_5_ to 1,5(PP)_2_-InsP_4_, or Kcs1, which converts InsP_6_ to 5PP-InsP_5_ and 1PP-InsP_5_ to 1,5(PP)_2_-InsP ^43, 44^ (Supplementary Fig. 1c,d). We found that deletion of either kinase reduced the interaction between wild type AtSPX1 and AtPHR1^226–360^ (Supplementary Fig. 1c). The interaction between the plant brassinosteroid receptor kinase BRI1 and the inhibitor protein BKI1, known to occur independently of PP-InsPs^45^, was not effected in either Δvip1vip1 or Δvip1kcs1 mutants (Supplementary Fig. 1c).

Using quantitative isothermal titration calorimetry (ITC) binding assays, we have previously determined dissociation constants (K*_D_*) for InsP_6_ and InsP_7_ binding to a OsSPX4 – OsPHR2 complex to be ∼50 and ∼7 µM, respectively^31^. A side-by-side comparison of InsP_7_ and InsP_8_ binding to OsSPX4 – OsPHR2 by ITC revealed dissociation constants of ∼7 and ∼3 µM, respectively (Fig. 1b). Taken together, the SPX – PHR interaction is mediated by PP-InsPs, with the *bona fide* Pi signaling molecule InsP_8_ being the preferred ligand *in vitro*.

### AtSPX1 interacts with a unique four-stranded coiled-coil domain in AtPHR1

We next mapped the SPX – PP-InsP binding site in AtPHR1 to a fragment (AtPHR1^280–353^), which comprises the CC domain and a 30 amino-acid spanning N-terminal extension, in yeast-2-hybrid experiments (Fig. 1a). We sought to crystallize an AtSPX1 – PP-InsP – AtPHR1 complex either in the pre- or absence of P1BS fragments. We obtained crystals of a putative AtSPX1 – InsP_8_ - AtPHR1^280-360^ complex diffracting to 2.4 Å resolution, and solved the structure by molecular replacement, using isolated SPX domain structures as search models^31^. Iterative cycles of model building and crystallographic refinement yielded, to our surprise, a well-refined model of AtPHR1^280–360^ only (see Methods). Analysis with the program PISA revealed the presence of a crystallographic tetramer in which four long α-helices fold into an unusual anti-parallel four-stranded coiled-coil (Fig. 1c). AtPHR1^280-360^ residues 292-356 and 310-357 are visible in the electron density maps from chain A and B, respectively. Residues 292-311 in chain A fold into a protruding loop region that harbors a small α-helix, and appear disordered in chain B (Fig. 1c). The anti-parallel ɑ-helices in AtPHR1 closely align with a root mean square deviation, (r.m.s.d.) of ∼ 0.5 Å comparing 45 corresponding C_ɑ_ atoms. Structural homology searches with the program DALI^46^ returned different coiled-coil structures, with a monomer of the tetrameric coiled-coil domain of the yeast transcription factor Ctp1 representing the closest hit (DALI Z-score 5.9, r.m.s.d. is ∼1 Å comparing 45 corresponding C_ɑ_ atoms) (Fig. 1d)^47^. However, no anti-parallel four-stranded coiled-coil domain with structural similarity to AtPHR1 was recovered, with, for example, the Ctp1 dimer-of-dimers domain having a very different configuration (Fig. 1e)^47^.

We next assessed the oligomeric state of AtPHR^280–360^ using size-exclusion chromatography coupled to right-angle light scattering (SEC-RALS) and determined an apparent molecular weight of ∼37.5 kDa, thus confirming that the isolated AtPHR1 CC forms tetramers in solution (theoretical molecular weight of the monomer is ∼9.5 kDa) (Fig. 2a). Two additional crystal structures of AtPHR1^280-360^ obtained in different crystal lattices all revealed highly similar tetrameric arrangements (Supplementary Fig. 2, Table 5).

**Fig. 2.**
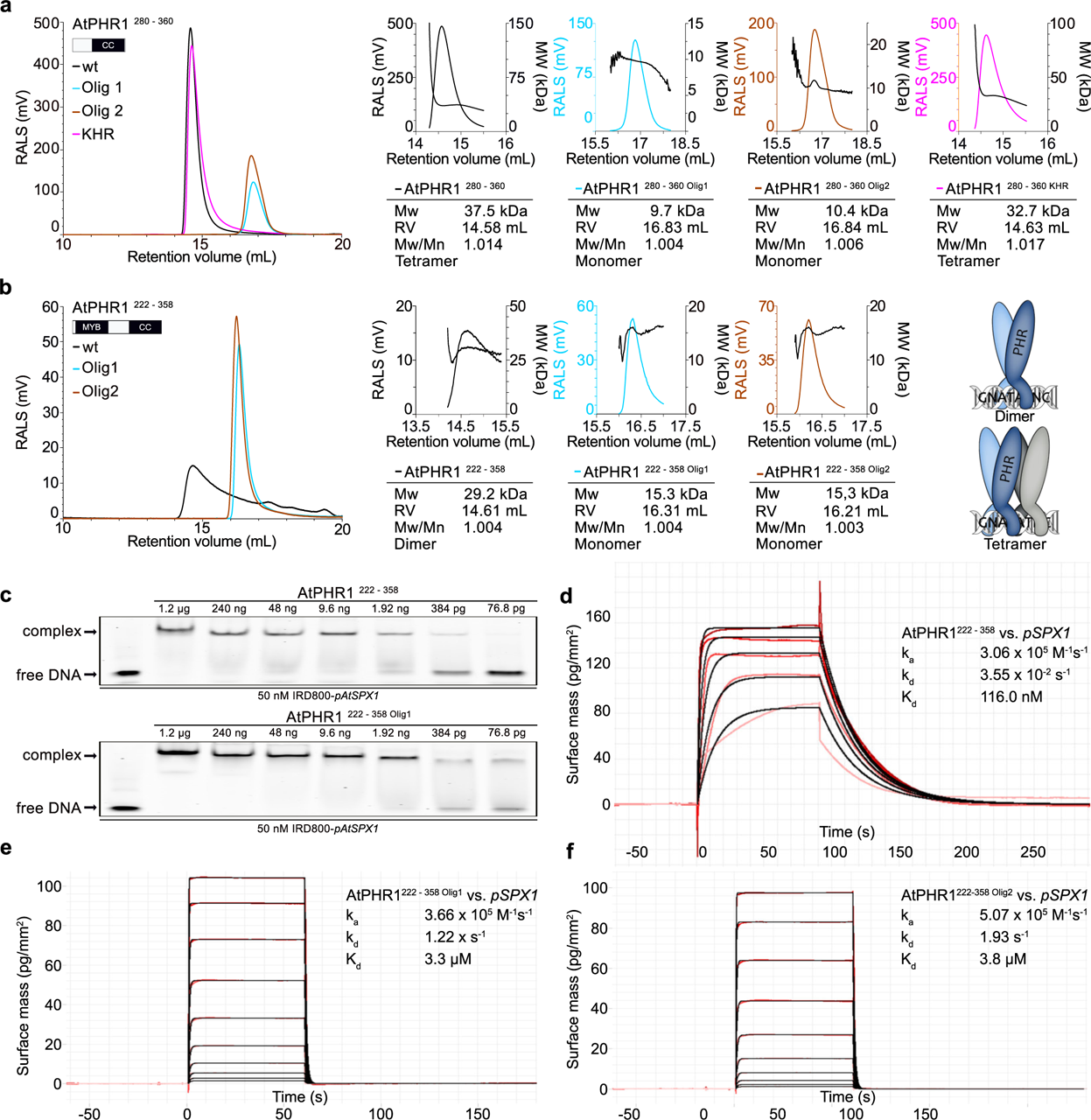
Mutations in the AtPHR1 CC domain impair oligomerisation and DNA binding. **a** Analytical size exclusion chromatography traces of wild type AtPHR1 CC (wt, black line), AtPHR1^280 – 360 Olig1^ (Olig 1, cyan), AtPHR1^280 – 360 Olig2^ (Olig 2, orange), and of AtPHR1^280 –360 KHR^ (KHR, magenta). The corresponding right-angle light scattering (RALS) traces are shown alongside, the molecular masses are depicted by a black line. Table summaries provide the molecular weight (Mw), retention volume (RV), dispersity (Mw/Mn), and the derived oligomeric state of the respective sample. **b** Analysis of AtPHR1 MYB CC (AtPHR1^222-358^) as described in **a**. **c** Qualitative comparison of the interaction of AtPHR1^222 – 358^ (upper panel) or AtPHR1^222 – 358 Olig1^ (lower panel) binding to IRD800-*pAtSPX1* in electrophoretic mobility shift assays. **d-f**, Quantitative comparison of the interaction of AtPHR1^222 – 358^, AtPHR1^222 – 358 Olig1^, or AtPHR1^222 – 358 Olig2^ with *pSPX1* by grating-coupled interferometry (GCI). Sensorgrams show raw data (red lines) and their respective fits (black lines). Table summaries provide the derived association rate (k_a_), the dissociation rate (k_d_) and the dissociation constant (K_d_).

### Mutations in the CC domain abolish AtPHR1 oligomerization and DNA binding *in vitro*

It has been recently reported that a AtPHR1^208 - 360^ fragment, which contains both the MYB and the CC domains fused to a maltose binding protein (MBP) tag, forms monomers in solution^48^. In contrast, our AtPHR1^280-360^ CC fragment is a tetramer (Fig. 2a). We thus purified untagged AtPHR1^222 – 358^ that comprises the CC and the MYB DNA-binding domains, and performed SEC-RALS experiments. We found that AtPHR1^222-358^ behaves as a dimer in solution (Fig. 2a,b; black traces), in agreement with earlier reports^2^. The observed oligomeric state differences between our AtPHR1^280-360^ (CC) and AtPHR1^222-358^ MYB-CC prompted us to investigate the dimer- and tetramerization interfaces in our AtPHR1 structures with the program PISA^49^. We found the dimerization (∼1,400 Å^2^ buried surface area) and the tetramerization (∼1,900 Å^2^ buried surface area) interfaces to be mainly formed by hydrophobic interactions (Supplementary Fig. 3a,b). Both interfaces are further stabilized by hydrogen bond interactions and several salt bridges (Supplementary Fig. 3a,b). Importantly, all contributing amino-acids represent sequence fingerprints of the plant unqiue MYB-CC transcription factor subfamily and are highly conserved among different plant species (Supplementary Fig. 3c). We identified residues specifically contributing to the formation of a CC dimer (Olig1: AtPHR1^L319^, AtPHR1^I333^, AtPHR1^L337^, shown in cyan in Fig. 2 and Supplementary Fig. 3) or tetramer (Olig2: AtPHR1^L317^, AtPHR1^L327^, AtPHR1^I341^, shown in dark-orange in Fig. 2 and Supplementary Fig. 3) in our different CC structures (Supplementary Table 5). We replaced these residues by asparagine to generate two triple mutants in AtPHR1^222 – 358^ and AtPHR1^280–360^, respectively. We found in SEC-RALS assays that both mutant combinations dissolved AtPHR1^280–360^ tetramers and AtPHR1^222–358^ dimers into stable monomers, respectively (Fig. 2a,b).

It has been recently reported that the AtPHR1 MYB DNA-binding domain associates with its target DNA as a dimer^42^. We thus studied the capacity of AtPHR1^222 – 358^ oligmerization mutants to interact with the P1BS in qualitative EMSA and quantitative grating coupled interferometry (GCI) assays. AtPHR1^222 – 358 Olig1^ could still interact with the P1BS in EMSAs indistinguishable from wild type (Fig. 2c). However, AtPHR1^222 – 358 Olig1^ and AtPHR1^222 – 358 Olig2^ bound a biotinylated P1BS immobilized on the GCI chip with ∼20-fold reduced affinity when compared to the wild type control (Fig. 2d-f). Together, our experiments suggest that PHR1 MYB-CC exists as a dimer in solution, and that disruption of its plant unique CC domain interface reduces the capacity of the transcription factor to bind its DNA recognition site.

### CC surface mutations abolish PHR – SPX interactions but do not interfere with DNA binding *in vitro*

We next sought to identify the binding site for SPX-InsP_8_ in the PHR CC domain. In our structures, a conserved set of basic residues maps to the surface of the four CC helices (shown in magenta in Fig. 3a, Supplementary Fig. 3c). A similar set of surface exposed basic residues has been previously found to form the binding site for PP-InsPs in various SPX receptors^31^. Mutation of AtPHR1^K325^, AtPHR1^H328^, AtPHR1^R335^, but not of AtPHR1^K308^, AtPHR1^R318^, AtPHR1^R340^ to alanine, disrupted the interaction of AtPHR1 with AtSPX1 in yeast (Fig. 3b). We next simultaneously mutated the residues corresponding to AtPHR1^K325^, AtPHR1^H328^, AtPHR1^R335^ to alanine in OsPHR2 (OsPHR2^KHR^). The mutant transcription factor showed no detectable binding to OsSPX4-InsP_7_ in quantitative ITC assays, but maintained the ability to bind the P1BS (Fig. 3c-d). In line with this, mutation of the KHR motif does not alter the oligomeric state of AtPHR1^280-360^ as concluded from SEC-RALS experiments (Fig. 2a, magenta traces). Taken together, three highly conserved basic residues located at the surface of the PHR CC domain are critical for the interaction with the PP-InsP bound SPX receptor (Supplementary Fig. 3c).

**Fig. 3.**
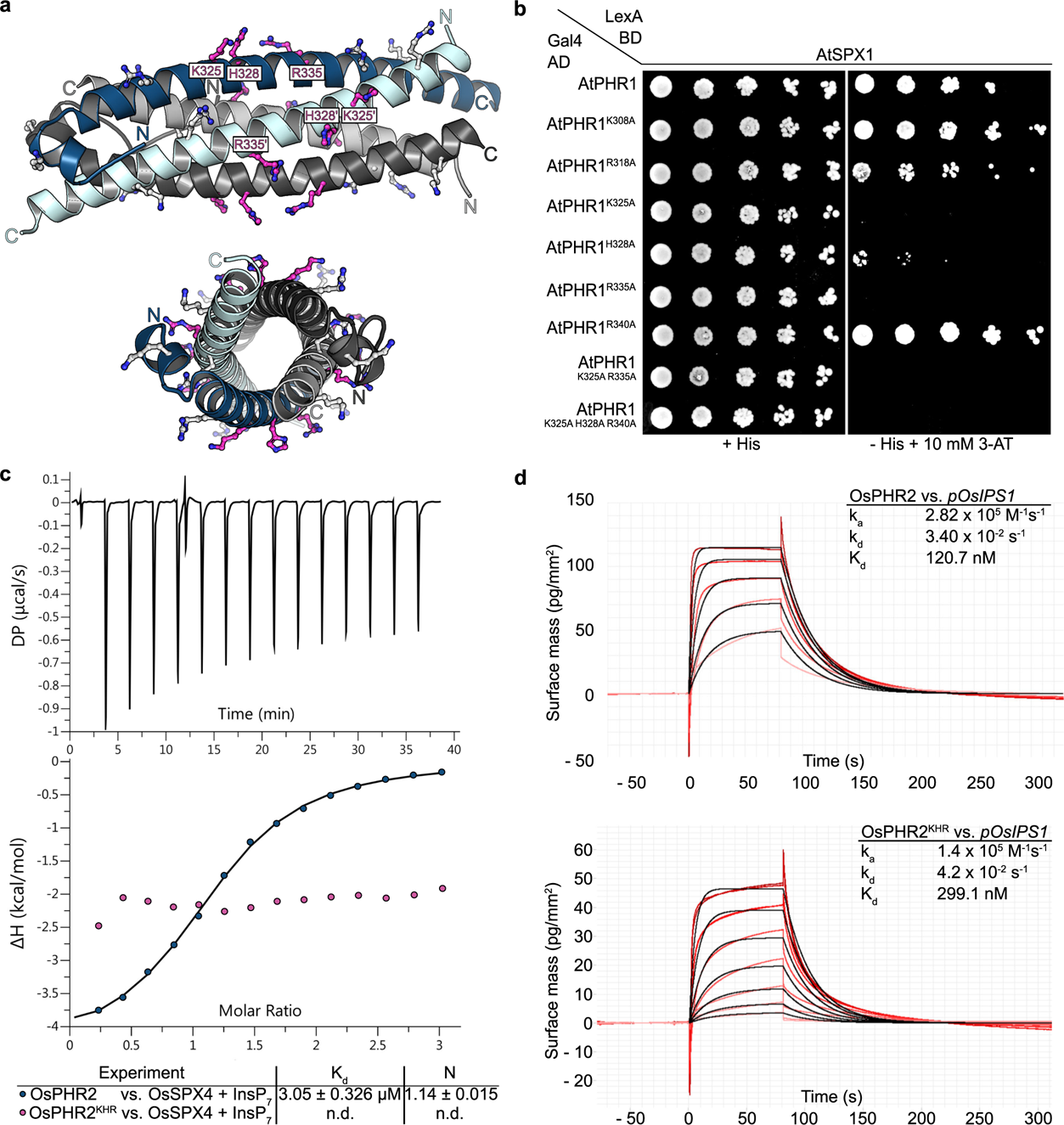
The KHR motif at the surface of the PHR CC is required for the interaction with SPX domains. **a** Ribbon diagram of the AtPHR1 CC domain with conserved basic residues located at the surface of the domain shown in bonds representation. The KHR motif (AtSPX1^K325^, AtSPX1^H328^, AtSPX1^R335^) is highlighted in magenta. **b** Mutational analysis of the basic residues in AtPHR1 CC. Yeast co-expressing AtPHR1^226 – 360^ variants in which surface exposed basic residues have been replaced with alanine fused to the Gal4-AD (prey) and AtSPX1 fused to the LexA-BD (bait) were grown on selective SD medium supplemented with histidine (+ His; co-transformation control) or lacking histidine and supplemented with 10 mM 3-amini-1,2,4-triazole (3-AT) (-His + 10 mM 3-AT; interaction assay) to identify residues required for interaction with AtSPX1 in yeast two-hybrid assays. Shown are serial dilutions from lift to right. **c** Isothermal titration calorimetry (ITC) assay of wild-type OsPHR2 and OsPHR2^KHR^ (300 µM) versus OsSPX4 (20 µM) - 5PP-InsP_5_ (100 µM). Raw heats per injection are shown in the top panel, the bottom panel represents the integrated heats of each injection, fitted to a one-site binding model (solid line). The insets show the dissociation constant (*K*_d_) and binding stoichiometry (N) (± fitting error, n.d. no detectable binding). **d** Quantitative comparison of the interaction of OsPHR2 (top panel) or OsPHR2^KHR^ (bottom panel) with *pOsIPS1* by GCI. Sensorgrams show raw data (red lines) and their respective fits (black lines). The insets show summarize association rates (k_a_), dissociation rates (k_d_) and the dissociation constant (K_d_) of the respective sample.

### Mutation of the AtPHR1 KHR motif impairs AtSPX1 binding and Pi homeostasis *in planta*

We next tested if mutation of the SPX binding site in AtPHR1 can modulate its function in Pi homeostasis in Arabidopsis. We expressed wild-type and point-mutant versions of AtPHR1 carrying an N-terminal FLAG tag under the control of its native promoter in a *phr1-3* loss-of-function mutant^9^. At seedling stage, we found that AtPHR1 single, double and triple point mutations complemented the previously characterized Pi deficiency phenotype of *phr1-3*^2, 9^. (Fig. 4a, Supplementary Fig. 4a). After transferring the seedlings to soil, variable growth phenotypes became apparent 21 days after germination (DAG) (Supplementary Fig 4b). From three independent lines per genotype, we selected one line each showing similar *AtPHR1* transcript levels for all experiments shown in Figure 4 (Fig. 4, Supplementary Fig. 5a). Comparing these lines, we found that AtPHR1^K325A,R335A^ double and AtPHR1^K325A,H328A,R335A^ triple mutants, but not the single mutants displayed severe growth phenotypes, with the triple mutant showing the strongest defects (Fig. 4a). We next determined cellular Pi levels in all independent lines and found that i) Pi levels are positively correlated with AtPHR1 expression levels (Supplementary Fig. 5a), that ii) all AtPHR1 mutant proteins tested accumulate Pi to significantly higher levels when compared to wild type and *phr1-3*, and that iii) the AtPHR1^K325A,H328A,R335A^ triple mutant displayed the highest Pi levels (Fig. 4b, Supplementary Fig. 5b-d). In line with this, PSI gene expression is misregulated in AtPHR1^K325A,R335A^ double and AtPHR1^K325A,H328A,R335A^ triple mutants (Fig. 4c). In co-immunoprecipitation assays in *Nicotiana benthamiana* and in Arabidopsis, we found the interaction of AtPHR1^K325A,H328A,R335A^ with AtSPX1 to be reduced when compared to wild-type AtPHR1 (Fig. 4d, Supplementary Fig. 6).

**Fig. 4.**
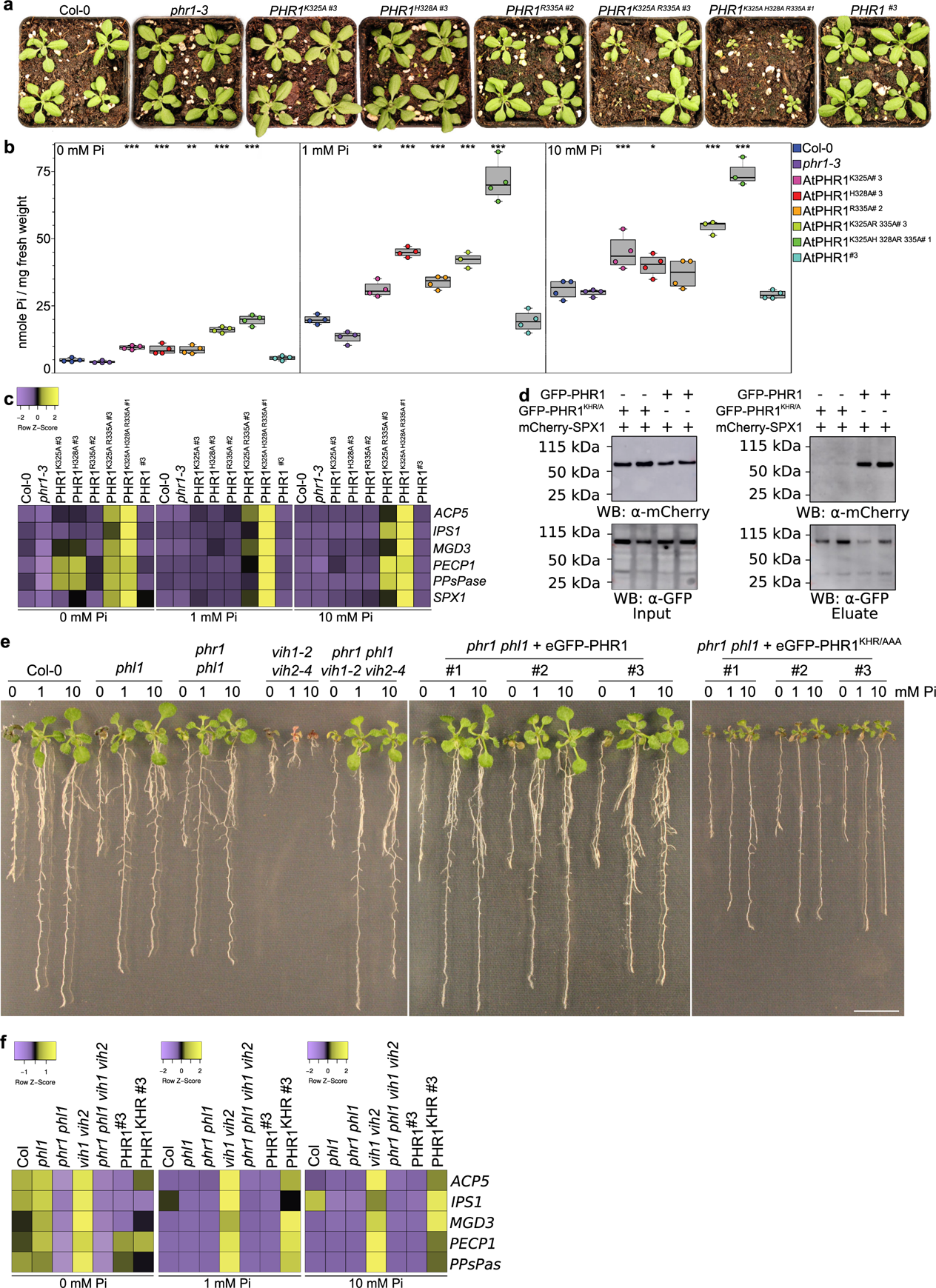
Mutation of the AtPHR1 KHR motif impairs interaction with AtSPX1 and Pi homeostasis *in planta*. **a** Growth phenotypes of Col-0 wild type, *phr1-3* of *phr1-3* complementation lines expressing FLAG-AtPHR1, FLAG-AtPHR1^K325A^, FLAG-AtPHR1^H328A^, FLAG-AtPHR1^R335A^, FLAG-AtPHR1^K325A R335A^, and FLAG-AtPHR1^K325A H328A R335A^ under the control of the *AtPHR1* promoter at 21 DAG grown in Pi sufficient conditions. One representative line per complementation construct is shown (specified by a #), additional lines are shown in Supplementary Fig. 4. **b** Pi content of the lines shown in a. Seedlings were germinated and grown on vertical ^1/2^MS plates for 8 d, transferred to ^1/2^MS plates supplemented with either 0 mM, 1 mM or 10 mM Pi and grown for additional 7 d. For each line, four plants were measured in technical duplicates. Pi contents of all lines can be found in Supplementary Fig. 4. **c** Heat maps of PSI marker gene (*ACP5, IPS1, MGD3, PECP1, PPsPase, SPX1*) expression analyses of the lines shown in **a**, represented as Z-scores. For each line, three biological replicates were analysed in technical triplicates by qRT-PCR. **d** Co-immunoprecipitation (Co-IP) experiment assessing the ability for immobilized GFP-AtPHR1 and GFP-AtPHR1^KHR/A^ to interact with mCherry-AtSPX1 in *N. benthamiana*. Input western blots are shown alongside. **e** Genetic interactions in the VIH-PHR signalling pathway. Col-0 wild type and the indicated mutant seedlings were grown on ^1/2^MS plates for 7 DAG, transferred to ^1/2^MS plates supplemented with either 0 mM, 1 mM or 10 mM Pi and grown for additional 7 d. For complementation analyses, wild-type AtPHR1 or AtPHR1^KHR/AAA^ was expressed as an N-terminal eGFP fusion protein under the control of the *AtPHR1* promoter. **f** Heat maps of PSI marker gene expression for the lines shown in **e**.

We next studied the genetic interaction between *PHR1* and *VIH1/2*. As previously reported, the severe phenotypes of *vih1-2 vih2-4* seedlings are partially rescued in the *phr1 phl1 vih1-2 vih2-4* quadruple mutant, suggesting that VIH1/2 generated InsP_8_ regulates the activity of PHR1 and PHL1 by promoting the binding of SPX receptors^37,32^. We performed the orthogonal genetic experiment, by complementing the *phr1 phl1* mutant with AtPHR1^KHR/AAA^ expressed under the control of the AtPHR1 promoter and carrying a N-terminal eGFP tag (Fig. 4e, see Methods). The complemented lines displayed intermediate growth phenotypes and constitutive PSI gene expression (Fig. 4e,f). Thus, SPX-InsP_8_ mediated regulation of PHR1 and PHL1 has to be considered one of several PP-InsP regulated processes affected in the *vih1-2 vih2-4* mutant. Together, our *in vivo* experiments reveal that SPX receptors interact with the CC domain of AtPHR1 via the surface exposed Lys325, His328 and Arg335, and that this interaction negatively regulates PHR activity and Pi starvation responses.

## Discussion

PHR transcription factors have been early on recognized as central components of the PSR in green algae and in plants, directly regulating the expression of PSI genes^1, 2, 12^. In Arabidopsis and in rice *spx* mutants of then unknown function also showed altered PSI gene expression^22, 24^. This genetic interaction was later substantiated by demonstrating that stand-alone plant SPX proteins can interact with PHR orthologs from Arabidopsis and rice^19, 20, 25^. The biochemical characterization of SPX domains as cellular receptors for PP-InsPs and the genetic identification of VIH kinases as master regulators of PSR in plants suggested that PP-InsPs, and specifically InsP_8_ mediates the interaction of SPX proteins with PHRs in response to changing nutrient conditions^31, 32, 37^.

Our quantitative DNA binding assays demonstrate that AtPHR1 MYB-CC binds the P1BS from the *AtSPX1* promoter with high affinity (K*_D_* ∼ 0.2 µM), in agreement with previously reported binding constants for different MYB-CC constructs (K*_D_* ∼0.01 – 0.1µM)^28, 42^. Different oligomeric states have been reported for various PHR MYB-CC constructs^2, 28^. Our AtPHR1 MYB-CC construct behaves as a dimer in solution, consistent with the recently reported crystal structure of the AtPHR1 MYB – DNA complex, and with earlier reports^2, 42^.

We could not assess the oligomeric state of full-length AtPHR1, due to rapid degradation of the recombinant protein. In yeast-2-hybrid assays, we found that AtSPX1-4 all are able to interact with AtPHR1 (Fig. 1a, Supplementary Fig. 1a). We mapped their conserved interaction surface to the plant-unique CC motif of PHRs (Fig. 1a). Crystal structures of this fragment reveal an unusual, four-stranded anti-parallel coiled-coil domain (Fig. 1c). Given the fact, that AtPHR1 MYB-CC forms dimers in solution (Fig. 2b), we cannot exclude the possibility that the CC tetramers represent crystal packing artifacts. However, we did observe identical CC tetramers in three independent crystal lattices (Supplementary Fig. 2) and in solution (Fig. 2a). The residues contributing to the dimer and to the tetramer interfaces are highly conserved among all plant MYB-CC transcription factors (Supplementary Fig. 3). Mutation of either interface blocks AtPHR1 oligomerization *in vitro* (Fig. 2a,b), and reduces DNA binding (Fig 2d-f). An attractive hypothesis would thus be that AtPHR1 binds its target promoter as a dimer, but can potentially form hetero-tetramers with other MYB-CC type transcription factors sharing the conserved, plant-unique CC structure and sequence (Supplementary Fig. 3). Notably, PHR1 PHL1 heteromers have been previously described^12^.

We found that SPX – PHR complex formation is mediated by endogenous PP-InsPs in yeast cells, as deletion of the yeast PP-InsP kinases Vip1 and Kcs1 abolished the interaction, and mutation of the PP-InsP binding surface in AtSPX1 interfered with AtPHR1 binding (Supplementary Fig. 1b,c). In line with this, SPX – PHR complexes are found dissociated in *vih1 vih2* mutant plants^32^. It is of note that the observed differences in binding affinity for InsP_7_ and InsP_8_ to SPX – PHR *in vitro* (Fig. 1b)^32^ cannot fully rationalize the apparent preference for InsP_8_ *in vivo*^32, 37^. We identified the binding surface for SPX-InsP_8_ by locating a set of highly conserved basic residues exposed at the surface of the CC domain (Fig. 3a, Supplementary Fig. 3). Mutation of this KHR motif did not strongly impact the ability of isolated OsPHR2 to bind *pOsIPS1 in vitro* (Fig. 3d), but disrupted the interaction of AtPHR1 with AtSPX1 in yeast (Fig. 3b). The corresponding mutations in OsPHR2 had a similar effect on the interaction with OsSPX4 in quantitative ITC assays (Fig. 3c). Expression of AtPHR1^KHR/AAA^ in the *phr1-3* mutant resulted in Pi hyperaccumlation phenotypes and constitutive PSI gene expression in Arabidopsis (Fig. 4a-c). The intermediate growth phenotypes of *vih1 vih2 phr1 phl1* mutants complemented with AtPHR1^KHR/AAA^ clearly suggests that PP-InsPs do not only regulate the activity of PHR1 and PHL1 in plants, but likely the function of other (SPX domain-containing) proteins^31^ (Fig. 4e). Notably, binding of AtPHR1^KHR^ to AtSPX1 was reduced in co-immunoprecipitation assays when compared to wild type AtSPX1 (Fig. 4d, Supplementary Fig. 6). Thus, our and previous finding suggest that InsP_8_ can act as a molecular glue, promoting the association of SPX receptors and PHR transcription factors. The interaction surface is likely formed by the previously characterized SPX basic surface patch^31^, and by the newly identified basic surface area in PHR CC, harboring the conserved KHR motif (Fig. 3a).

It is of note that addition of AtSPX1 to AtPHR1 has been previously demonstrated to reduce AtPHR1’s ability to bind to P1BS in the presence of InsP_6_ ^28^. We could not quantify these interactions in ITC or GCI binding assays, as PHR CC formation is much preferred over SPX-InsP_8_ binding at the protein concentrations required in these assays. We speculate that InsP_8_ bound SPX proteins can bind to the basic residues we identified in the PHR CC domain to control the oligomeric state and hence the promoter binding capacity of PHRs. As these residues are conserved among all plant MYB-CC proteins this may suggest that transcription factors outside the PHR sub-family may be regulated by SPX domains and PP-InsPs, possibly rationalizing the severe phenotypes of *vih1 vih2* mutant plants (Fig. 4e, Supplementary Fig. 3). The recent findings that VIH kinases and PHRs act together in plant PSR and that SPX-PHR complexes are dissociated in *vih1 vih2* mutants further suggest that InsP_8_ is the *bona fide* signaling molecule promoting the association between SPX domains and MYB-CCs^32, 37^.

Repressive SPX – PHR complexes consequently form only under Pi sufficient conditions, where InsP_8_ levels are high^32, 37^. Under Pi starvation, when InsP_8_ levels are reduced, SPX - PHR complexes dissociate, enabling the transcription factors to acquire the oligomeric state required for high affinity promoter binding. The physiological and mechanistic investigation of this central process may, in the long term, contribute to the development of Pi starvation resilient crops. This could in turn sustain the use of the essential and non-renewable resource rock phosphate, which is currently consumed at an alarming scale.

## Methods

### Molecular cloning, constructs and primers

For a detailed description of the cloning strategies, constructs and primers used in this study, please refer to Supplementary Table 2 and Table 3a-c.

### Generation of stable transgenic *A. thaliana* lines

All stable transgenic *A. thaliana* lines are listed in Supplementary Table 1. Constructs were introduced into *Agrobacterium tumefaciens* strain pGV2260 and *A. thaliana* plants were transformed via floral dipping^50^. Transformants were identified by mCherry fluorescence with a Zeiss Axio Zoom.V16 stereo microscope (mRFP filter) and a HXP200C illuminator. Homozygous T3 lines have been identified for complementation lines expressing FLAG-AtPHR1, FLAG-AtPHR1^K325A^, FLAG-AtPHR1^H328A^, FLAG-AtPHR1^R335A^ under the control of the native *AtPHR1* promoter. For complementation lines expressing FLAG-AtPHR1^K325A R335A^ and FLAG-AtPHR1^K325A H328A R335A^ under the control of the native *AtPHR1* promoter, T2 lines were used throughout, and homozygous and heterozygous transformants were selected for all experiments by mCherry fluorescence as described above. T3 homozygous lines expressing eGFP-PHR1 under the control of the *AtPHR1* promoter were identified by their Hygromycin resistance. PHR1 was amplified from Arabidopsis cDNA and introduced into pH7m34GW binary vector. Point mutations were introduced by site-directed mutagenesis (primers are listed in Supplementary Table 3f).

### Yeast-two hybrid experiments

(Screen) AtSPX1^1–252^ was used as a bait and screened against an *A. thaliana* seedling cDNA library by Hybrigenics Services. AtSPX1^1–252^ was cloned into the pB29 vector providing a C-terminal LexA-DNA binding domain (BD) and transformed into yeast strain L40αGal4 (MATa). Prey genes were cloned into the pP6 vector providing a N-terminal Gal4 activation domain (AD) and transformed into yeast strain YHGX13 (MATα). After mating haploid bait and prey strains, positive interactions were detected by growth on histidine deficient medium. (Yeast strains and media) For all experiments, either the diploid TATA strain (Hybrigenics Services) or the haploid L40 strain was used (Supplementary Table 2). Cells were routinely maintained on YPAD plates (20 g/L glucose, 20 g/L bacto-peptone, 10 g/L yeast extract, 0.04 g/L adenine hemisulfate, 20 g/L agar). Experiments were performed on synthetic dropout (SD) plates (6.7 g/L yeast nitrogen base with adenine hemisulfate and without leucine, tryptophan and histidine, 20 g/L glucose, 20 g/L agar) supplemented with 0.076 g/L histidine or 10 mM 3-amino,1,2,4-triazole (3-AT).

(Yeast transformation) One yeast colony was resuspended in 500 µL sterile H_2_O, plated on YPAD plates and grown for two days until the whole plate was covered with yeast. Yeast cells were then resuspended in 50 mL YPAD liquid medium and the OD_600nm_ was determined (2*10^6^ cells/mL were used for one transformations). Cells were centrifuged at 3,000 xg and 4°C for 5 min, resuspended in 25 mL TE buffer, centrifuged again at 3,000 xg and 4°C for 5 min, resuspended in 2 mL LiAc/TE buffer, centrifuged at 16,000 xg and RT for 15 s, and finally resuspended in 50µL/transformation LiAc/TE buffer. The transformation mix (0.5 µg bait plasmid, 0.5 µg prey plasmid, 10 µL ssDNA (10 mg/mL), 50 µL yeast cells, 345 µL 40% (w/v) PEG3350 in LiAC/TE) was prepared and incubated at 30°C for 45 minutes, followed by incubation at 42°C for 30 min. Finally, yeast cells were centrifuged at 6,500 xg and RT for 15s, resuspended in TE buffer, plated on SD plates lacking leucine and tryptophan and incubated at 30°C for 3 days. (Yeast spotting dilution assay) Positive transformants were selected on SD plates without tryptophan and leucine, and incubated at 30°C for three days. Cells were counted, washed in sterile water and spotted in 5 times dilution (5000, 1000, 200, 40, 8 cells) on SD plates without either tryptophan and leucine, or tryptophan, leucine and histidine supplemented with 10 mM 3-AT. Plates were incubated at 30°C for 3 days.

### Protein expression and purification

(AtPHR1, OsPHR2 and OsSPX4) AtPHR1^280–360 wt/KHR/Olig1/Olig2^ and AtPHR1^222-358 wt/Olig1/Olig2^ were cloned into the pMH-HT protein expression vector, providing a N-terminal 6xHis affinity tag with a tobacco etch virus (TEV) protease recognition site. OsPHR2^1-426 wt/KHR^ was cloned into the pMH-HSgb1T protein expression vector, providing a N-terminal 8xHis-Strep-gb1 affinity tag with a TEV cleavage site. OsSPX2^1-321^ was cloned into the pMH-HSsumo protein expression vector, providing a N-terminal 8xHis-Strep-Sumo affinity tag. All constructs were transformed into *E. coli* BL21 (DE3) RIL cells. For recombinant protein expression, cells were grown at 37°C in terrific broth (TB) medium to an OD_600nm_ of 0.6. After reducing the temperature to 18°C, protein expression was induced with 0.3 mM isopropyl β-D-galactoside (IPTG) for 16 h. Cells were centrifuged at 2,200 rpm and 4°C for 1 h, resuspended in lysis buffer (50 mM Tris-HCl pH 7.8, 500 mM NaCl, 0.1% (v/v) IGEPAL, 1mM MgCl, 2mM β-mercaptoethanol), snap-frozen in liquid nitrogen and stored at −80°C. For protein preparation, cells were thawed, supplemented with cOmplete^TM^ EDTA-free protease inhibitor cocktail (Roch), DNaseI, and lysozyme, and disrupted using a sonicator. Cell lysates were centrifuged at 7,000 xg and 4°C for 1 h, sterile filtered, supplemented with 20 mM imidazole and loaded onto a 5 mL HisTrap HP Ni^2+^ affinity column (GE Healthcare). After washing with several column volumes (CV) of lysis buffer supplemented with 20 mM imidazole, high salt buffer (50 mM Tris-HCl pH 7.8, 1M NaCl, 2 mM β-mercaptoethanol), and high phosphate buffer (200 mM K_2_HPO_4_/KH_2_PO_4_ pH 7.8, 2 mM β-mercaptoethanol), proteins were eluted in a gradient from 20 to 500 mM imidazole in lysis buffer. The purified proteins were cleaved by TEV or Sumo protease overnight at 4°C (1:100 ratio) and dialyzed against lysis buffer for PHR1^222-358 wt/Olig1/Olig2^ and PHR1^280-360 wt/KHR/Olig1/Olig2^ fragments, against modified lysis buffer (25 mM Tris-HCl pH 7.8, 300 mM NaCl, 0.1% (v/v) IGEPAL, 1mM MgCl, 2mM β-mercaptoethanol) for OsPHR2^1-426 wt/KHR^, and against modified anion exchange buffer (20 mM Tris-HCl pH 6.5, 500 mM NaCl) for OsSPX4^1-321^. PHR1^280-360 wt/KHR/Olig1/Olig2^ and OsPHR2^1-426 wt/KHR^ were subjected to a second Ni^2+^ affinity purification in either lysis buffer or modified lysis buffer, respectively, and the flow-throughs were collected and concentrated. PHR1^222-358 wt/Olig1/Olig2^ were subjected to cation exchange (50 mM Hepes pH 7.5, 50 – 1000 mM NaCl), and OsSPX4^1-321^ was subjected to anion exchange (20 mM Tris-HCl, 50 – 1000 mM NaCl). Fractions corresponding to the respective proteins were pooled and concentrated. All proteins were loaded onto a HiLoad Superdex 75pg HR26/60 column (GE healthcare), pre-equilibrated in gel filtration buffer A (20 mM Tris/HCl pH 7.5, 300 mM NaCl, 0.5 mM TCEP) for PHR1^222-358 wt/Olig1/Olig2^, or in gel filtration buffer B (20 mM Tris/HCl pH 7, 200 mM NaCl, 0.5 mM TCEP) for the remaining proteins. Fractions containing the respective proteins were pooled and concentrated. Purified and concentrated protein was immediately used for further experiments or snap-frozen in liquid nitrogen and stored at −80°C.

(AtPHR1^280-360^ used for crystallization) The AtPHR1^280-360^ fragment was cloned into the pMH-HS-Sumo protein expression vector, providing a N-terminal 8xHis-StrepII tandem affinity tag and a Sumo fusion protein. The construct was transformed into *E. coli* BL21 (DE3) RIL cells. For recombinant protein expression, cells were grown at 37°C in terrific broth (TB) medium to an oD_600nm_ of 0.6. After reducing the temperature to 16°C, protein expression was induced with 0.3 mM isopropyl β-D-galactoside (IPTG) for 16 h. Cells were centrifuged for 20 min at 4,000 g and 4°C, then the cell pellet was washed with PBS, snap-frozen in liquid nitrogen and stored at −80°C. For protein complex purification, the cell pellet was thawed and mixed with twice the amount of cells expressing a His-Strep-MBP-AtSPX1^1-251^ fusion protein, which provides a N-terminal, TEV-cleavable maltose binding protein (MBP). Lysis buffer (200 mM KP_i_ pH 7.8, 2 β-ME) supplemented with 0.1% (v/v) IGEPAL, 1 mM MgCl_2_, 10 mM imidazole, 500 units TurboNuclease (BioVision), 2 tablets Protease Inhibitor Cocktail (Roche) was added and cells were disrupted using an EmulsiFlex-C3 (Avestin). Cell lysates were centrifuged at 7,000 g and 4°C for 1 h. The cleared supernatant was sterile filtered and loaded onto a 5 mL HisTrap HP Ni^2+^ affinity column (GE Healthcare). After washing with several column volumes of lysis buffer, the protein was eluted with 250 mM imidazole in lysis buffer. The purified His-Strep-Sumo-AtPHR1/His-Strep-MBP-AtSPX1 fusion proteins were cleaved by TEV and Sumo protease treatment overnight at 4°C while dialyzing in a buffer containing 200 mM KP_i_ pH 7.8, 100 mM NaCl, 2 β-ME. 10 mM imidazole were added to the cleaved protein sample and a second Ni^2+^ affinity step was performed in order to remove the cleaved-off His-Strep-Sumo/MBP fusion tags as well as the 6xHis-tagged Sumo and TEV proteases. The flow-through was concentrated and loaded onto a HiLoad Superdex 75pg HR16/60 column (GE healthcare), pre-equilibrated in gel filtration buffer (20 mM Tris/HCl pH 7.8, 250 mM NaCl, 2.5 mM InsP_6_, 0.5 mM TCEP). Fractions containing both, the AtSPX1^1-251^ and AtPHR1^280-360^ proteins, were pooled and concentrated. A second size exclusion chromatography step was performed using a HiLoad Superdex 200pg HR26/60 column (GE healthcare) and the same gel filtration buffer as above. Purified and concentrated protein was immediately used for further experiments or snap-frozen in liquid nitrogen and stored at −80°C.

### Isothermal titration calorimetry (ITC)

All ITC experiments were performed at 25°C using a MicroCal PEAQ-ITC system (Malvern Panalytical) equipped with a 200 μl sample cell and a 40 μl injection syringe. 5PP-InsPl sample cell and a 40 μl sample cell and a 40 μl injection syringe. 5PP-InsPl injection syringe. 5PP-InsP_5_ (InsP_7_) and 1,5(PP)_2_-InsP_4_ (InsP_8_) were produced as described^51^. All proteins were dialysed against ITC buffer (20 mM HEPES pH 7.0, 200 mM NaCl) and PP-InsP ligands were diluted in ITC buffer prior to all measurements. A typical titration consisted of 15 injections, the protein concentrations in the syringe and in the cell are provided in the respective figure legend. Data were analyzed using the MicroCal PEAQ-ITC analysis software (v1.21).

### Crystallization and data collection

Two hexagonal crystal forms developed in sitting drops consisting of 0.2 µL protein at a concentration of 12 mg/mL and 0.2 µL reservoir solution (0.1 M phosphate citrate pH 4.2, 0.2 M NaCl, 20% (w/v) PEG 8000). Crystals were cryo-protected by adding reservoir solution containing 10% (v/v) ethylene glycol directly to the drop and subsequently snap-frozen in liquid nitrogen. A third, tetragonal crystal form developed in 0.1 M Bis-Tri pH 6.5, 0.1 M NaCl, 1.5 M (NH_4_)_2_SO_4_. Crystals were cryo-protected by serial transfer into reservoir solution supplemented with 15% (v/v) glycerol and snap-frozen in liquid nitrogen. Crystal forms 1, 2 and 3 diffracted to ∼2.4, ∼2.5 Å and ∼1.9 Å resolution, respectively. Data were collected at beam-line PXIII of the Swiss Light Source (SLS), Villigen, Switzerland. Data processing and scaling was done in XDS^52^.

### Crystallographic structure solution and refinement

The AtPHR1^280-360^ structure was solved by molecular replacement using the previously described SPX^CtGde1^ (PDB-ID:5IJJ) core helices as search model in calculations with the program PHASER^53^. The structure was completed in iterative cycles of manual model building in COOT^54^ and restrained refinement in phenix.refine^55^ or Refmac5^56^. Residues 280-294 and residues 278-280 appear disordered in the final model. Quality of the structural model was assessed by using MolProbity^57^, refinement statistics are shown in Supplementary Table 5. Structure visualization was done with PyMOL (Molecular Graphics System, Version 1.7, Schrödinger, LLC) and ChimeraX^58^. The structure of AtSPX1 was modeled using the program SWISS-MODEL^59^ and the SPX^HsXPR1^ domain structure of the human phosphate exporter as template (PDB-ID:5IJH, GMQE score ∼ 0.49, QMEAN4 score ∼ −2.27, 29.5% sequence identity^60^). Conserved PP-InsP binding residues in AtSPX1 were determined by aligning sequences with previously described SPX domains^31^ using the program T-coffee^61^.

### Right angle light scattering (RALS)

The oligomeric state of AtPHR1 variants was analyzed by size-exclusion chromatography paired with a refractive-index detector using an OMNISEC RESOLVE/REVEAL combined system (Malvern). Instrument calibration was performed with a BSA standard (Thermo Scientific Albumin Standard). Samples of 50 µL containing 2 – 10 mg/mL AtPHR1 (wild type AtPHR1^280–360^, AtPHR1^280-360 Olig1^, AtPHR1^280-360 Olig2^, AtPHR1^280-360 KHR^, wild type AtPHR1^222-358^, AtPHR1^222-358 Olig1^, AtPHR1^222-358 Olig2^) in OMNISEC buffer (20 mM Hepes pH 7.5, 150 mM NaCl) were separated on a Superdex 200 increase 10/300 GL column (GE Healthcare) at a column temperature of 25°C and a flow rate of 0.7 ml min^−1^. Data were analyzed using the OMNISEC software (v10.41).

### DNA oligonucleotide annealing

DNA oligonucleotides were dissolved in annealing buffer (10 mM HEPES-NaOH pH 8.0, 50 mM NaCl, 0.1 mM EDTA). Equal volumes of the equimolar DNA oligonucleotides were mixed and incubated in a heat block for 5 min at 95 °C. Subsequently, DNA oligonucleotides were cooled down to room temperature for 90 min. Double-stranded DNA oligonucleotides were aliquoted and stored at −20°C.

### Electrophoretic mobility shift assay (EMSA)

5 % Mini-PROTEAN TBE precast gels (Bio-Rad) have been pre-electrophoresed in 0.5x TBE buffer for 60 minutes at 70 V. Reactions mixes have been prepared following the Odyssey^®^ Infrared EMSA kit manual (LI-COR) without the use of optional components, including 50 nM of IRDye800 end-labeled oligos (refer to Supplementary Table 4a; Metabion), and a 1:5 dilution series of wild type AtPHR1^222-358^ or AtPHR1^222-358 Olig1^ (1.2 µg to 76.8 pg). Reaction mixes have been incubated for 30 minutes at room temperature in the dark, and 2 µL of 10x Orange Loading Dye (LI-COR) have been added to each sample prior to loading on a 5% TBE gel. Gels have been electrophoresed until orange dye migrated to the bottom of the gel (∼ 1 h) at 70 V in the dark. Gels have been scanned with the 800 nm channel of an Odyssey imaging system (LI-COR).

### Grating coupled interferometry (GCI)

All GCI experiments were performed at 4°C using a Creoptix WAVE system (Creoptix sensors) with 4PCP WAVE chips (Creoptix sensors). Chips were conditioned with borate buffer (100 mM sodium borate pH 9.0, 1 M NaCl) and subsequently neutravidin was immobilized on the chip surface via standard amine-coupling: activation (1:1 mix of 400 mM *N*-(3-dimethylaminopropyl)-*N*′-ethylcarbodiimide hydrochloride, and 100 mM *N*-hydroxysuccinimide); immobilization (30 µg ml^−1^ of neutravidin in 10 mM sodium acetate, pH 5.0); passivation (5% BSA in 10 mM sodium acetate, pH 5.0); quenching (1 M ethanolamine, pH 8.0). Biotinylated oligos (Supplementary Table 4b; Metabion) were captured on the chip. Analytes were injected in a 1:2 dilution series starting from 4 µM (AtPHR1^222-358^), 20 µM (AtPHR1^223-358 Olig1^, AtPHR1^223-358 Olig2^), or 10 µM (OsPHR2, OsPHR2^KHR^) in GCI buffer (for OsPHR2: 20 mM HEPES pH 7.9, 200 mM NaCl; for AtPHR1: 20 mM HEPES pH 7.5, 300 mM NaCl). Blank injections every 4^th^ cycle were used for double referencing and a dimethylsulfoxide (DMSO) calibration curve (0%, 0.5%, 1%, 1.5%, 2%) for bulk correction. Data were corrected and analysed using the Creoptix WAVE control software (corrections applied: X and Y offset; DMSO calibration; double referencing; refractive index correction), and a one-to-one binding model was used to fit all experiments.

### Plant material, seed sterilization and plant growth conditions

All *A. thaliana* plants used in this study were of the Columbia (Col-0) ecotype. Seeds of the T-DNA insertion lines *phr1-3* (SALK_067629) and phl1 (SAIL_731_B09) were obtained from the European Arabidopsis Stock Center. Homozygous *phr1-3* and *phl1* lines were identified by PCR using T-DNA left and right border primers paired with gene-specific sense and antisense primers (Supplementary Table 3d). The *phr1 phl1* double mutant was kindly provided by Yves Poirier (University of Lausanne, Switzerland), *vih1-2 vih2-4* double and *phr1 phl1 vih1-2 vih2-4* quadruple mutants have been reported previously^37^. Seeds were surface sterilized by incubation in 70% (v/v) CH_3_-CH_2_-OH for 10 minutes, followed by incubation in 0.5% (v/v) sodium hypochlorite for 10 minutes, and subsequently washed four times in sterile H_2_O. Seeds were placed on full half-strength Murashige-Skoog plates^63^ containing 1 (w/v) % sucrose and 0.8 (w/v) % agar (^1/2^MS plates), and stratified for 2 to 3 d at 4°C in the dark prior to transfer into a growth cabinet. Plants were grown on vertical ^1/2^MS plates at 22°C under long day conditions (16 h light – 8 h dark) for 8 to 11 d.

### Western blot

Proteins were transferred to nitrocellulose membrane (GE Healthcare, Amersham^TM^ Highbond^TM^-ECL) via wet western blotting at 4°C and 30 V overnight. Membranes were blocked in TBS-Tween (0.1%) - Milk (5%) for one hour at room temperature. For mCherry detection, membranes were incubated overnight with an anti-mCherry antibody (ab167453, dilution 1:2000; Abcam) followed by one hour incubation with an anti-rabbit-HRP antibody (dilution 1:10000, Calbiochem). For GFP detection, membranes were incubated overnight with an anti-GFP-HRP antibody (130-091-833, dilution 1:1000, Miltenyi Biotec). For FLAG detection, membranes were incubated overnight with an anti-FLAG-HRP antibody (A8692, dilution 1:1000, Sigma). Antibodies were diluted in TBS-Tween (0.05%) - Milk (2.5%). Membranes were detected with SuperSignal™ West Femto Maximum Sensitivity Substrate (34095, Thermo Scientific™) and subsequently stained with Ponceau stain (0.1% (w/v) Ponceau S in 5% (v/v) acetic acid).

### Determination of cellular Pi concentrations

To determine cellular Pi concentration at seedling stage, plants were transferred from ^1/2^MS plates to -Pi ^1/2^MS plates containing 1 (w/v) % sucrose and 0.8 (w/v) % agarose supplemented with either 0 mM, 1 mM, or 10 mM Pi (KH_2_PO_4_/K_2_HPO_4_; pH 5.7) at 7 DAG, and grown at 22 °C under long day conditions. At 14 DAG, seedlings were weighted and harvested into 1.5 mL tubes containing 500 µL nanopure H_2_O. Samples were frozen at −80°C overnight, thawed at 80°C for ten minutes, refrozen at −80°C, incubated at 80°C and 1,400 rpm for one hour and briefly centrifuged to sediment plant tissue. Pi content was measured by the colorimetric molybdate assay^64^.

### RNA analyses

At 14 DAG, 50 - 150 mg seedlings were harvested in 2 ml Eppendorf tubes containing two metal beads each, shock-frozen in liquid nitrogen and ground in a tissue lyzer (MM400, Retsch). RNA extraction was performed using the ReliaPrep RNA Tissue Miniprep System (Promega) including in column DNase I treatment to remove genomic DNA. First strand cDNA synthesis was performed from 1 µg – 2.5 µg of total RNA using Superscript II RT (Invitrogen) with oligo(dT) primers. qRT-PCR was performed in 10 µL reactions containing 1x SYBR-Green fluorescent stain (Applied Biosystems) and measured using a 7900HT Fast Real Time PCR-System (Applied Biosystems). qRT-PCR programme: 2’-95°C; 40 x (30’’-95°C; 30’’-60°C; 20’’-72°C); melting curve 95°C – 60°C – 95°C. A primer list can be found in Supplementary Table 3e. Expression levels of target genes were normalized against the housekeeping gene *Actin2*. For every genotype, three biological replicates were analyzed in technical triplicates.

### Transient transformation of *Nicotiana benthamiana*

For each construct, 4 ml of *A. tumefaciens* strain pGV2260 suspension culture were grown overnight at 28°C. Cells were collected by centrifugation at 700 xg for 15 mins and resuspended in transformation buffer (10 mM MgCl_2_, 10 mM MES pH 5.6, 150 µM acetosyringone). Cell density was measured and set to a final OD_600_ of 0.5 for SPX1 and PHR1, and to 0.1 for the silencing suppressor P19. Suspension cultures were incubated for two hours in the dark at room temperature and subsequently mixed at a volume ratio of 1:1:1 (SPX1:PHR1:P19). *N. benthamiana* leaves were infiltrated using a 0.5-ml syringe and 3 leaf disks (d = 1 cm) per sample were harvested after 3 d, snap-frozen in liquid nitrogen, and stored at – 80°C.

### Co-immunoprecipitation

For co-immunoprecipitation experiments with proteins transiently expressed in *N. benthamiana*, samples were ground in liquid nitrogen with plastic mortars and proteins were extracted with 600 µL of homogenisation buffer (50 mM Tris-HCl pH 7.5, 150 mM NaCl, 0.25% Triton X-100, 5% (v/v) glycerol, 1 mM PMSF, cOmplete^TM^ EDTA-free protease inhibitor cocktail (Roche). Samples were incubated at 4°C for 10 min with gently rotation and subsequently centrifuged at 16,000 xg and 4°C for 15 min. Supernatants were transferred to fresh tubes and further centrifuged at 16,000 xg and 4°C for 15 min. Supernatants were transferred to fresh tubes, while 50 µL of each supernatant were taken and mixed with 10 µL 6x SDS sample buffer (input), the remaining supernatants were mixed with 50 µL magnetic µMACS anti-GFP beads (Miltenyi Biotec) and incubated at 4°C for 2 h with gently rotation. MACS columns (Miltenyi Biotec) were used with a µMACS Separator (Milteyi Biotec). MACS columns were washed with 200 µL of homogenisation buffer and samples were loaded. Columns were washed either four times with 200 µL of homogenisation buffer and once with wash buffer 2 (Miltenyi Biotec), or three times with 200 µL of homogenisation buffer, three times with wash buffer 1 (Miltenyi Biotec) and once with wash buffer 2. Columns were incubated with 20 µL preheated elution buffer (Miltenyi Biotec) for 5 min at room temperature. 50 µL of elution buffer were added and eluates were recovered. Inputs and eluates were boiled for 5 min at 95°C prior and separated on 9 % SDS-PAGE gels. Co-immunoprecipitation experiments for proteins stably or natively expressed in *A. thaliana*, were performed as previously described^65^.

### Statistics

Simultaneous inference was used throughout to limit the false positive decision rate in these randomized one- or two-way layouts. Designs with technical replicates were analysed using a mixed effect model. Normal distributed variance homogeneous errors were assumed when appropriate, otherwise a modified variance estimator allowing group-specific variances was used^66^. Multiple comparisons of several genotypes vs. wild type (Col-0) shown in Fig. 4b, and Supplementary Fig. 5a-d were performed as described^67^ (*, p < 0.5; **, p < 0.05).

### Data availability

Data supporting the findings of this manuscript are available from the corresponding authors upon reasonable request. A reporting summary for this article is available as a Supplementary Information file. The source data underlying the qPCR and Pi level measurements are provided as Source Data files. Coordinates and structure factors have been deposited in the Protein Data Bank (PDB) with accession codes 6TO5 (form1, https://doi.org/10.2210/pdb6TO5/pdb), 6TO9 (form2, https://doi.org/10.2210/pdb6TO9/pdb) and 6TOC (form 3, https://doi.org/10.2210/pdb6TOC/pdb). The associated X-ray diffraction images and data processing files have been deposited at http://zenodo.org with DOIs 10.5281/ zenodo.3570698 (form1), 10.5281/zenodo.3570977 (form 2) and 10.5281/zenodo.3571040 (form 3).

## Acknowledgements

This work was supported by European Research Council Consolidator Grant 818696/INSPIRE (to MH), by Swiss National Foundation Sinergia Grant CRSII5_170925 (to MH and DF), and by an HHMI International Research Scholar Award (to MH). MKR was supported by an EMBO long-term fellowship (ALTF-129-2017). We thank Irene Sabater for providing LI SPX1, and members of the Hothorn lab for critically reading the manuscript.

## Author contributions

MKR: Conceptualization, data curation, formal analysis, validation, investigation, visualization, methodology, and writing (original draft, review and editing). RW: Conceptualization, data curation, formal analysis, validation, investigation, methodology, and writing (review and editing). JZ: Investigation, methodology and writing (review and editing). RKH: Methodology. LB: Investigation and methodology. LAH: Software, formal analysis, methodology. DF: Resources, methodology, writing (review and editing). MH: Conceptualization, resources, data curation, formal analysis, supervision, funding acquisition, validation, investigation, visualization, methodology, project administration and writing (original draft, review, and editing).

---

**Supplementary Fig. 1.**
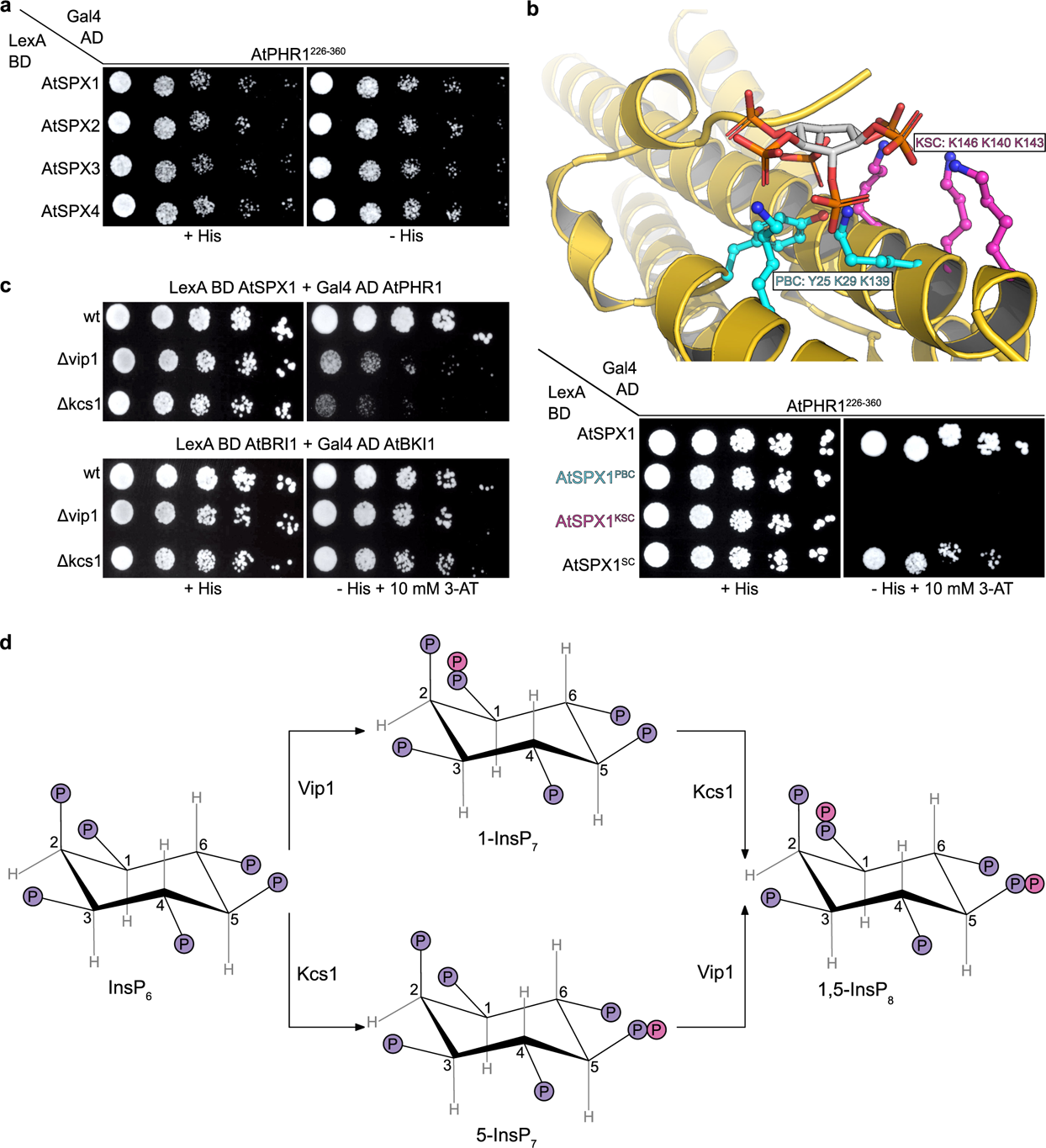
The AtPHR1 – AtSPX1 interaction in yeast is meditated by PP-InsPs. **a** Yeast-2-hybrid assay. Yeast co-expressing AtPHR1^226–360^ fused to the Gal4-AD (prey) and different AtSPX proteins fused to the LexA-BD (bait) were grown on selective SD medium supplemented with histidine (+ His; co-transformation control) or lacking histidine (-His; interaction assay). Shown are serial dilutions from left to right. **b** (Top panel) Homology model of an AtSPX1^1-182^-InsP_6_ complex. AtSPX1^1-182^ is shown as blue ribbon diagram and side chains involved in InsP_6_ binding are highlighted in green (PBC, phosphate binding cluster) and purple (KSC, lysine surface cluster) and depicted in bonds representation. The InsP_6_ ligand is shown in grey (in bonds representation). (Bottom panel) Yeast co-expressing AtPHR1^226-360^ fused to the Gal4-AD (prey) and different AtSPX1 versions mutated in residues involved in InsP_6_ binding, or a structural control mutant (SC^31^) fused to the LexA-BD (bait) were grown on selective SD medium supplemented with histidine (+ His; co-transformation control) or lacking histidine and supplemented with 10 mM 3-AT(-His + 3-AT; interaction assay) to investigate the importance of the PP-InsP binding surface in AtSPX1 for the AtSPX1 – AtPHR1 interaction in yeast. **c** Yeast knock-out strains for the PP-InsP biosynthesis enzymes Vip1 or Kcs1 co-expressing either AtPHR1^226-360^ fused to the Gal4-AD (prey) and AtSPX1 fused to the LexA-BD (bait) (upper panel), or AtBKI1 fused to the Gal4-AD (prey) and AtBRI1 fused to the LexA-BD (bait) (lower panel) were grown on selective SD medium supplemented with histidine (+ His; co-transformation control) or lacking histidine and supplemented with 10 mM 3-AT (-His + 3-AT; interaction assay) to investigate the importance of the availability of specific PP-InsPs for the AtSPX1-AtPHR1 interaction in yeast. **d** Schematic representation of the PP-InsP biosynthesis pathway in yeast.

**Supplementary Fig. 2.**
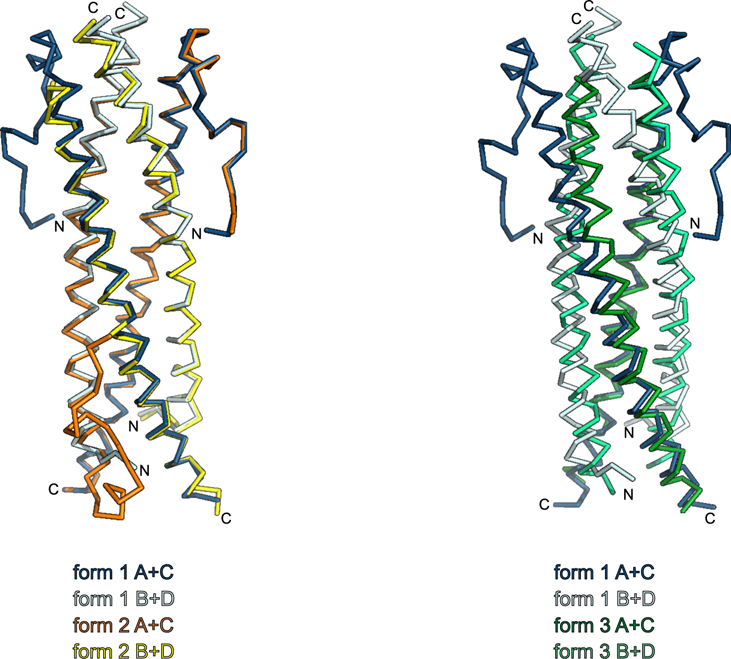
Three different AtPHR1 CC crystal structures all share the same tetrameric arrangement. Structural superposition (shown as C_ɑ_ traces) of the four-stranded anti-parallel coiled-coil domain of AtPHR1 from crystal forms 1-3.

**Supplementary Fig. 3.**
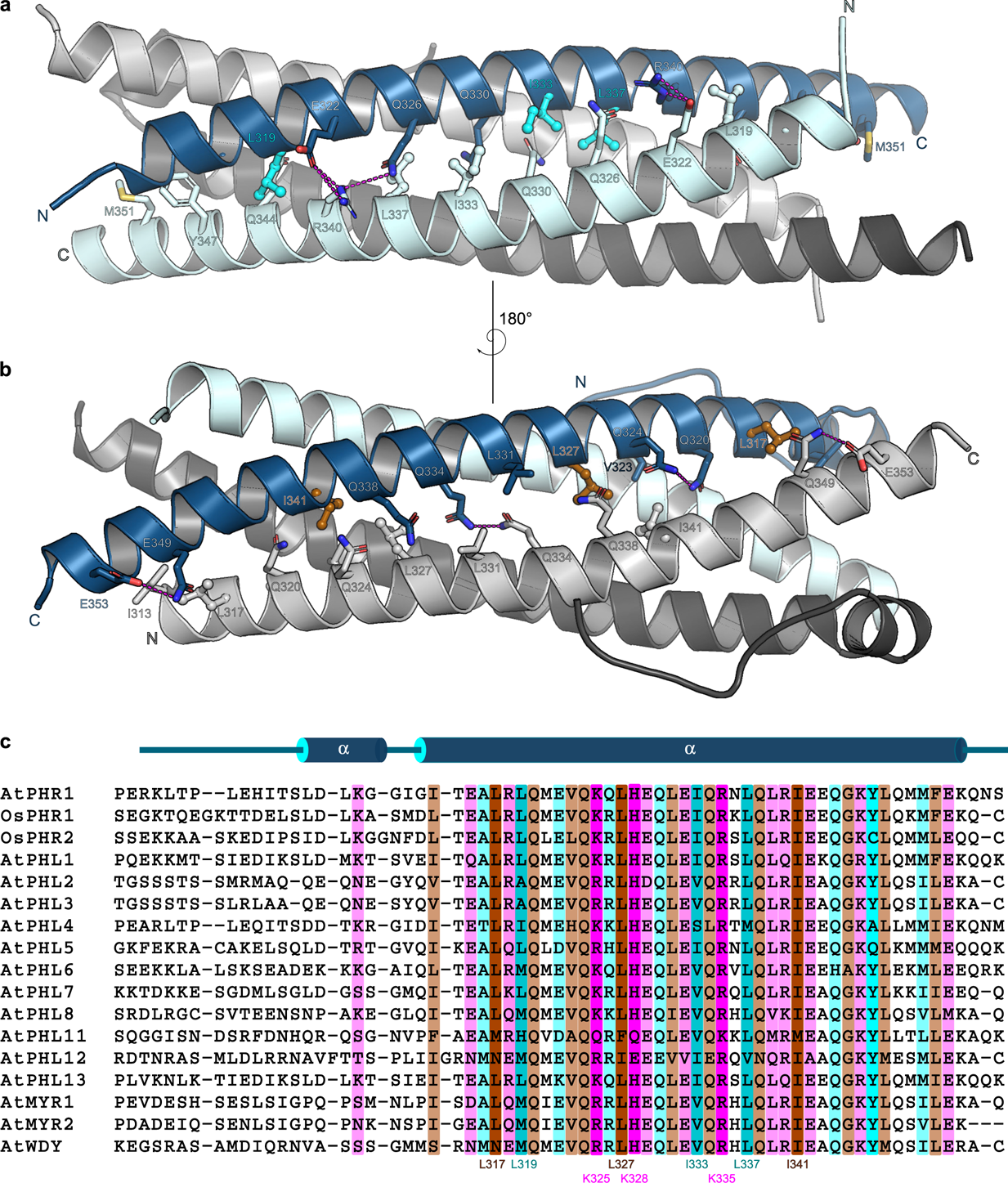
A conserved dimer- and tetramerization interface in plant MYB CC transcription factors. **a** Overview of the AtPHR1 CC dimerization interface. Shown is a ribbon diagram with selected residues contributing to the dimer interface shown in bonds representation. Hydrogen bonds are indicated as dotted lines, residues mutated in the Olig 1 mutants are highlighted in cyan. **b** Overview of the tetramerization interface, with residues mutated in Olig 2 depicted in gold. **c** Structure based sequence alignment of the CC domain of plant MYB CC domain and including a secondary structure assignment calculated with the program DSSP^73^. Residues contributing to the CC dimer interface are shown in blue and cyan, to the tetramerization interface in gold and brown, respectively. The conserved basic residues on the surface of the CC domain are highlighted in magenta.

**Supplementary Fig. 4.**
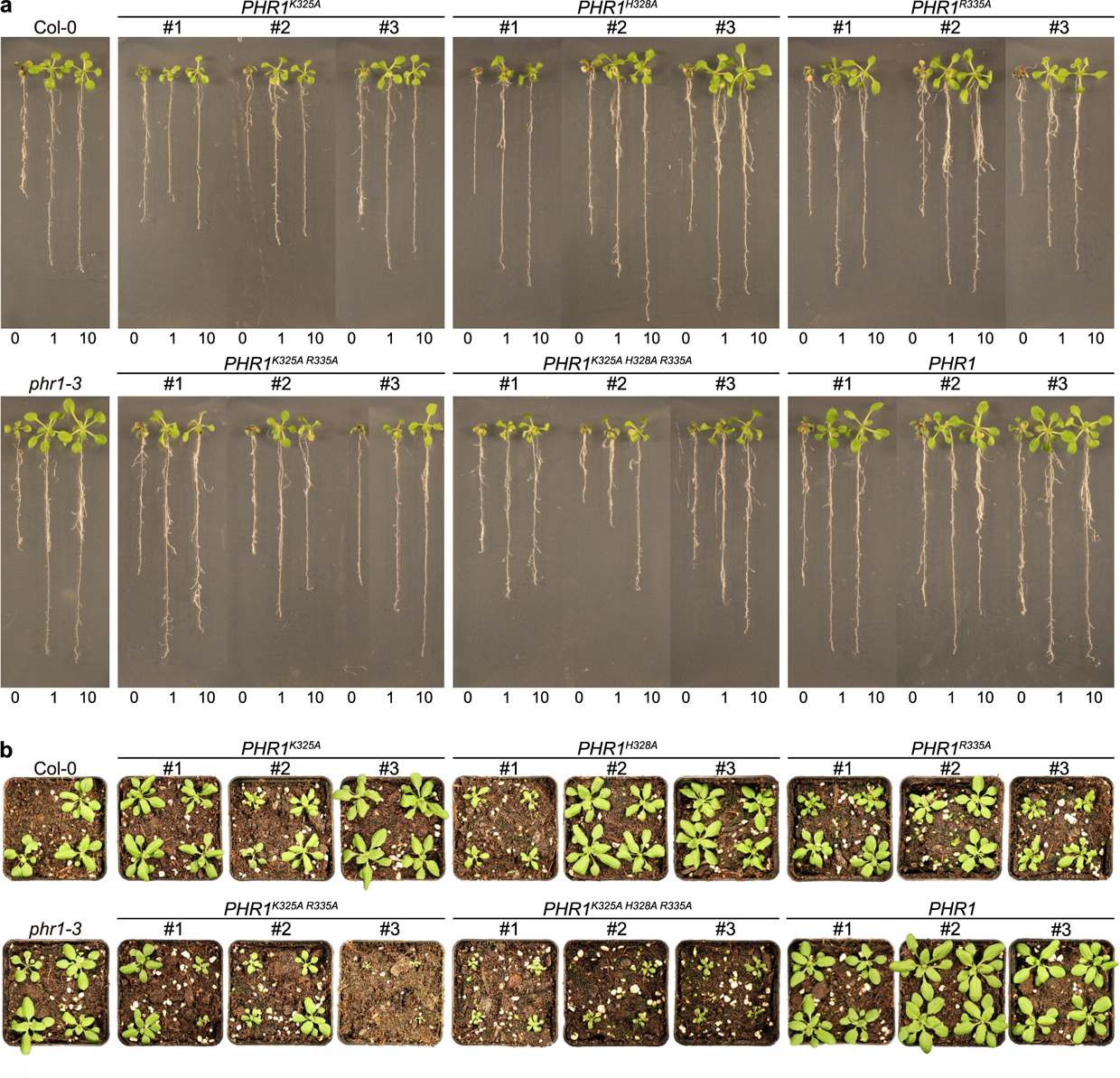
Growth phenotypes of AtPHR1 CC domain mutants that abolish interaction with AtSPX1 and impact Pi homeostasis. **a** Growth phenotype of Col-0 wild type, *phr1-3*, and seedlings of *phr1-3* complementation lines expressing FLAG-AtPHR1, FLAG-AtPHR1^K325A^, FLAG-AtPHR1^H328A^, FLAG-AtPHR1^R335A^, FLAG-AtPHR1^K325A R335A^, and FLAG-AtPHR1^K325A H328A R335A^ under the control of the *AtPHR1* promoter at 14 d after germination (DAG). Seedlings were germinated and grown on vertical ^1/2^MS plates for 8 d, transferred to ^1/2^MS plates supplemented with either 0 mM, 1 mM or 10 mM Pi and grown for additional 7 d. **b** Growth phenotypes of the lines in **a**, at 21 DAG. Seedlings were germinated and grown on vertical ^1/2^MS plates for eight days, transferred to soil and grown for additional 14 d.

**Supplementary Fig. 5.**
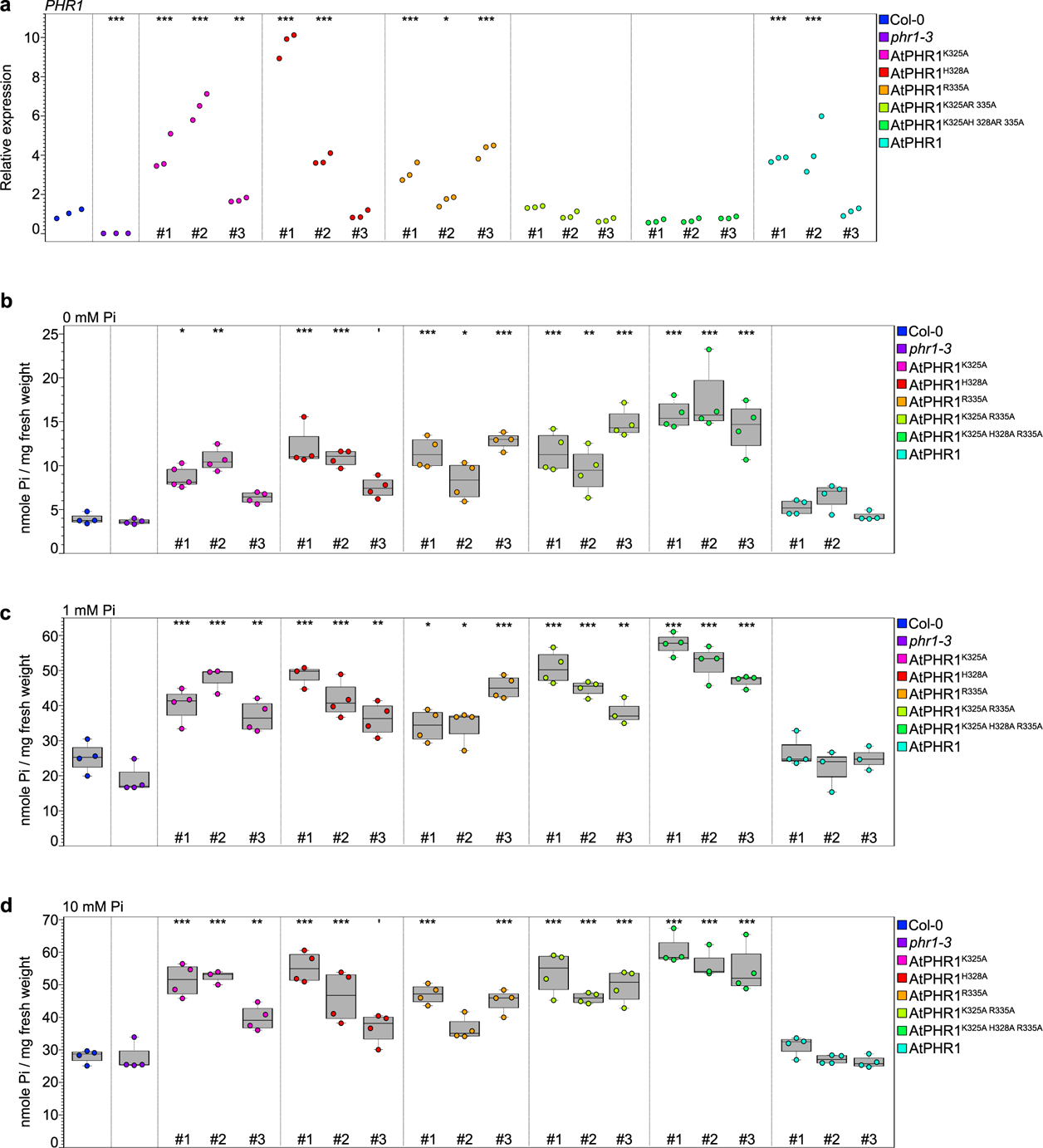
Mutations in the AtPHR1 KHR motif result in Pi hyperaccumulation. **a** Expression of *PHR1* in Col-0, *phr1-3*, and seedlings of *phr1-3* complementation lines expressing FLAG-AtPHR1, FLAG-AtPHR1^K325A^, FLAG-AtPHR1^H328A^, FLAG-AtPHR1^R335A^, FLAG-AtPHR1^K325A R335A^, and FLAG-AtPHR1^K325A H328A R335A^ under the control of the *AtPHR1* promoter relative to the housekeeping gene *Actin2* at 14 DAG. Seedlings were germinated and grown on vertical ^1/2^MS plates for 8 d, transferred to ^1/2^MS plates supplemented with 1 mM Pi and grown for additional 7 d. For each line, three biological replicates were analysed in technical triplicates by qRT-PCR. Stars indicate significant differences to Col-0 (Dunnett’s Test with Bonferroni correction; *, p < 0.05). **b-d** Pi content of Col-0 wild type, *phr1-3* seedlings and seedlings of *phr1-3* complementation lines described in **a**. Seedlings were germinated and grown on vertical ^1/2^MS plates for 8 d, transferred to ^1/2^MS plates supplemented with either 0 mM (b), 1 mM (c) or 10 mM (d) Pi and grown for additional 7 d. For each line, 4 plants were measured in technical duplicates. (*, p < 0.5; **, p < 0.05).

**Supplementary Fig. 6.**
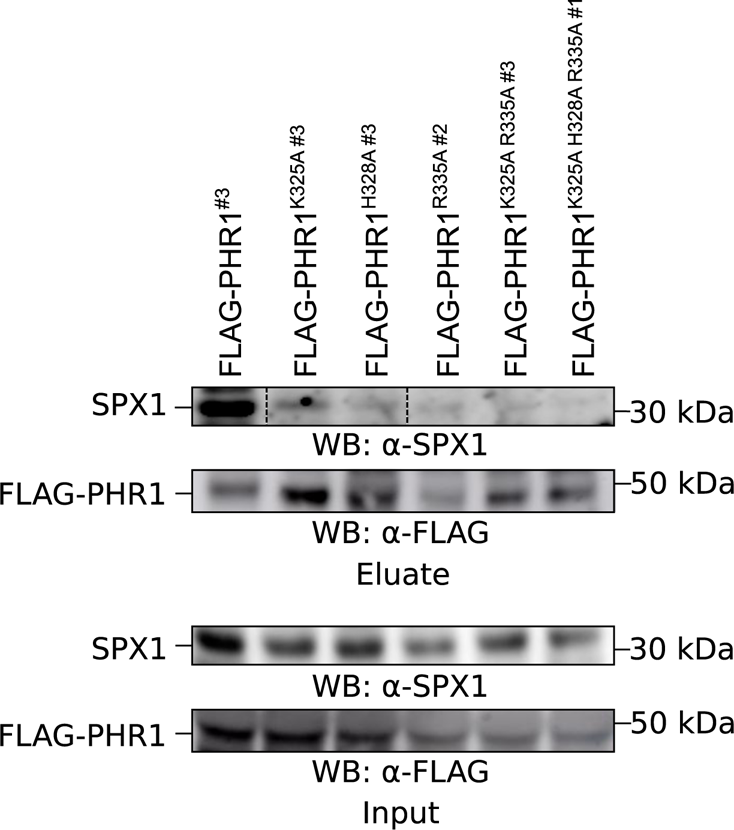
Mutation in the KHR motif reduces AtPHR1 binding to AtSPX1 in Arabidopsis. Co-immunoprecipitation experiments using FLAG-tagged wild type and mutant AtPHR1 variants stably expressed in Arabidopsis under the control of the *AtPHR1* promoter. Total protein was extracted from *phr1-3* complementation lines at 10 DAG. Seedlings were germinated and grown on vertical ^1/2^MS plates supplemented with 1 mM Pi. FLAG-tag fusions were affinity bound with magnetic FLAG-tag trap, and immunoprecipitation of FLAG-AtPHR1 was monitored by immunoblot with an anti-FLAG antibody. Co-enrichment of endogenous AtSPX1 was monitored by immunoblot with an anti-SPX1 antibody.

**Supplementary Table 1.**
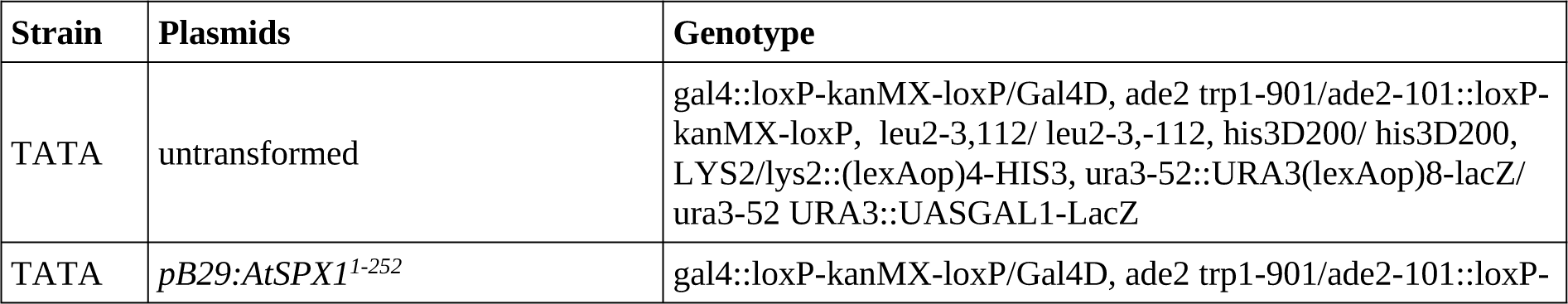
– Stable transgenic A. thaliana lines.

**Supplementary Table 2.**
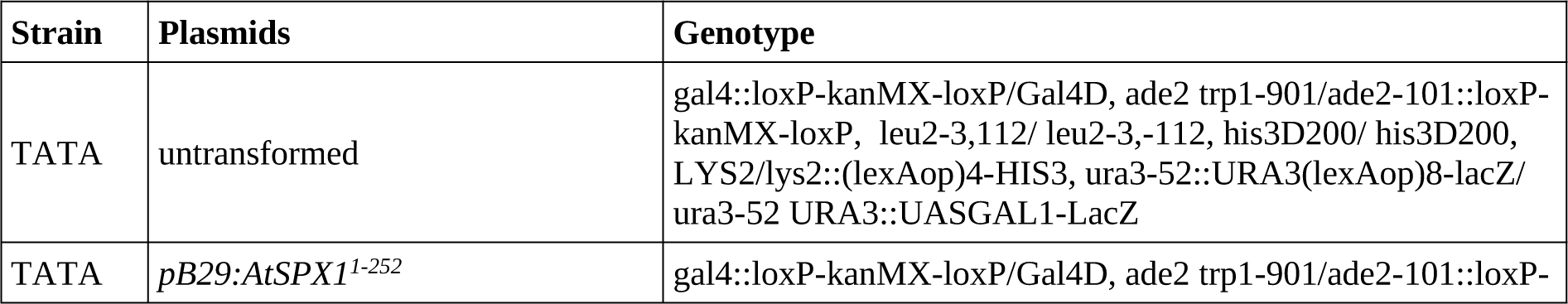
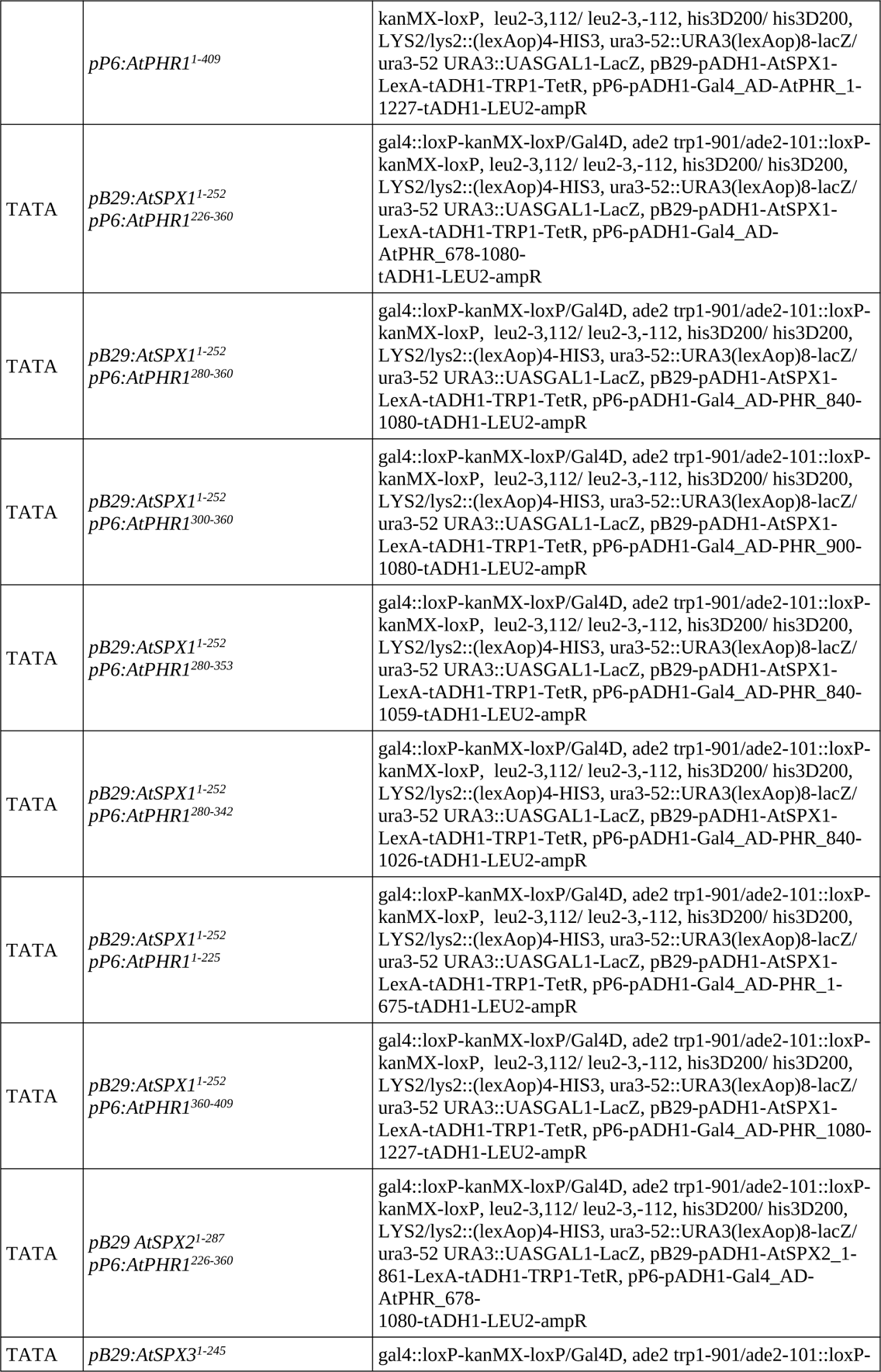
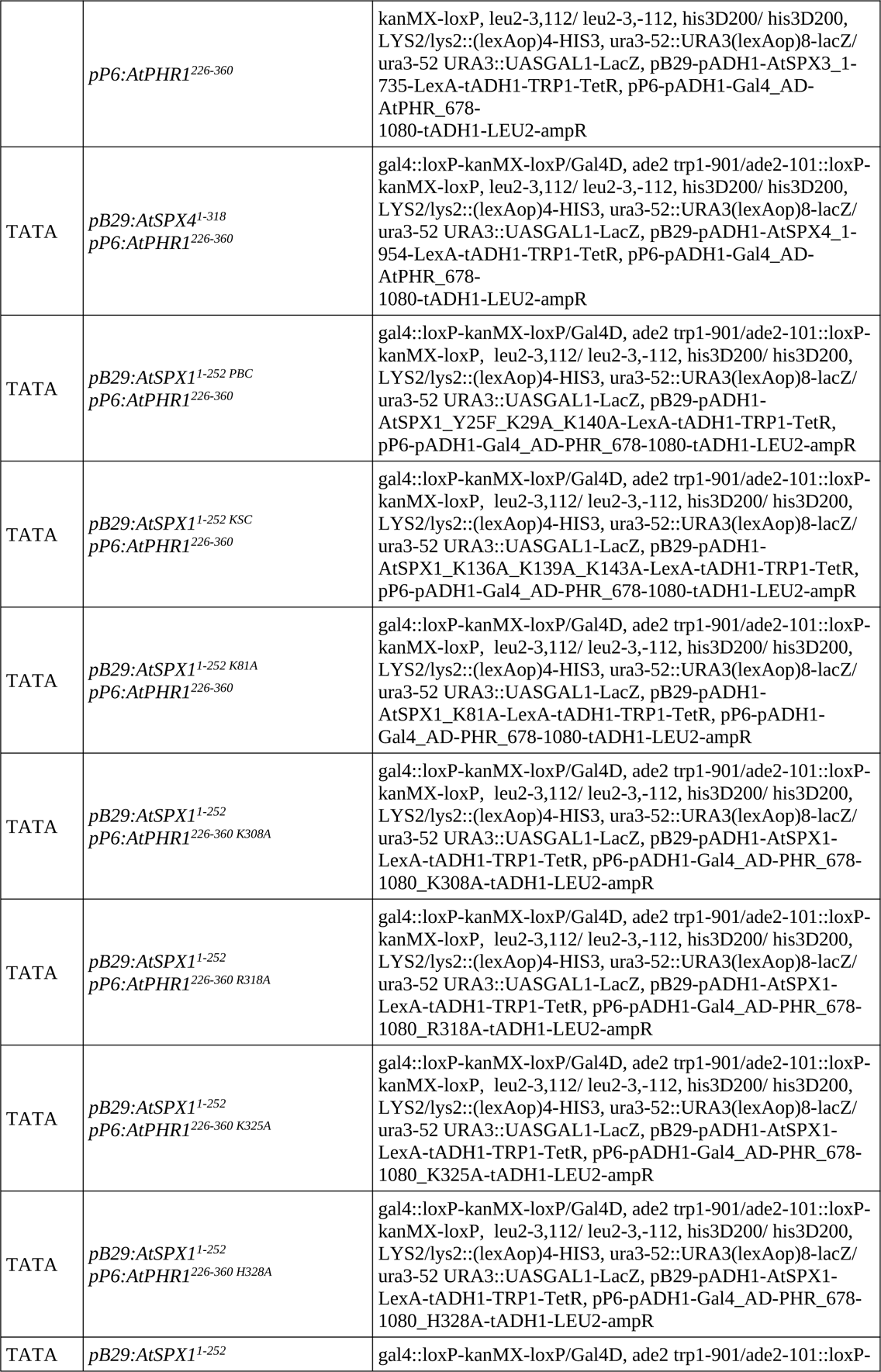
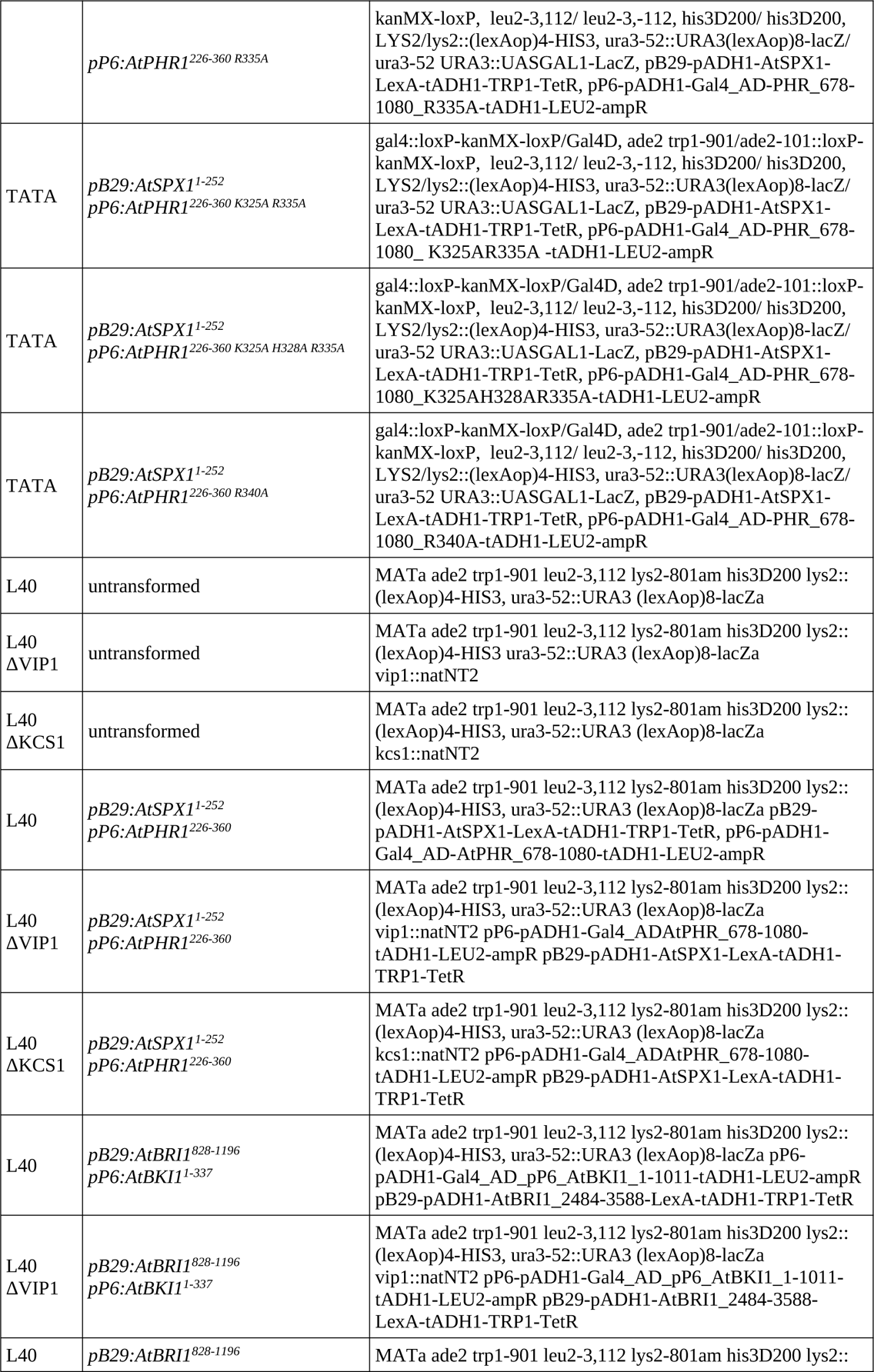
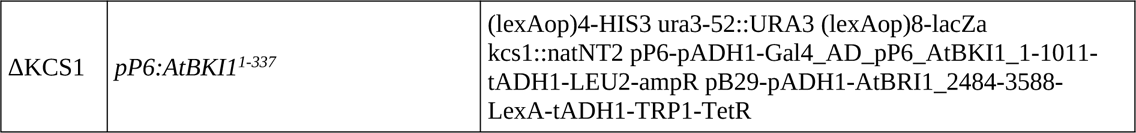
**– Yeast strains.** Plasmids for yeast transformation have been generated via Gibson cloning^68^. Mutations targeting AtPHR1 K325, H328 and R335 were introduced by site-directed mutagenesis PCR^69^.

**Supplementary Table 3.**
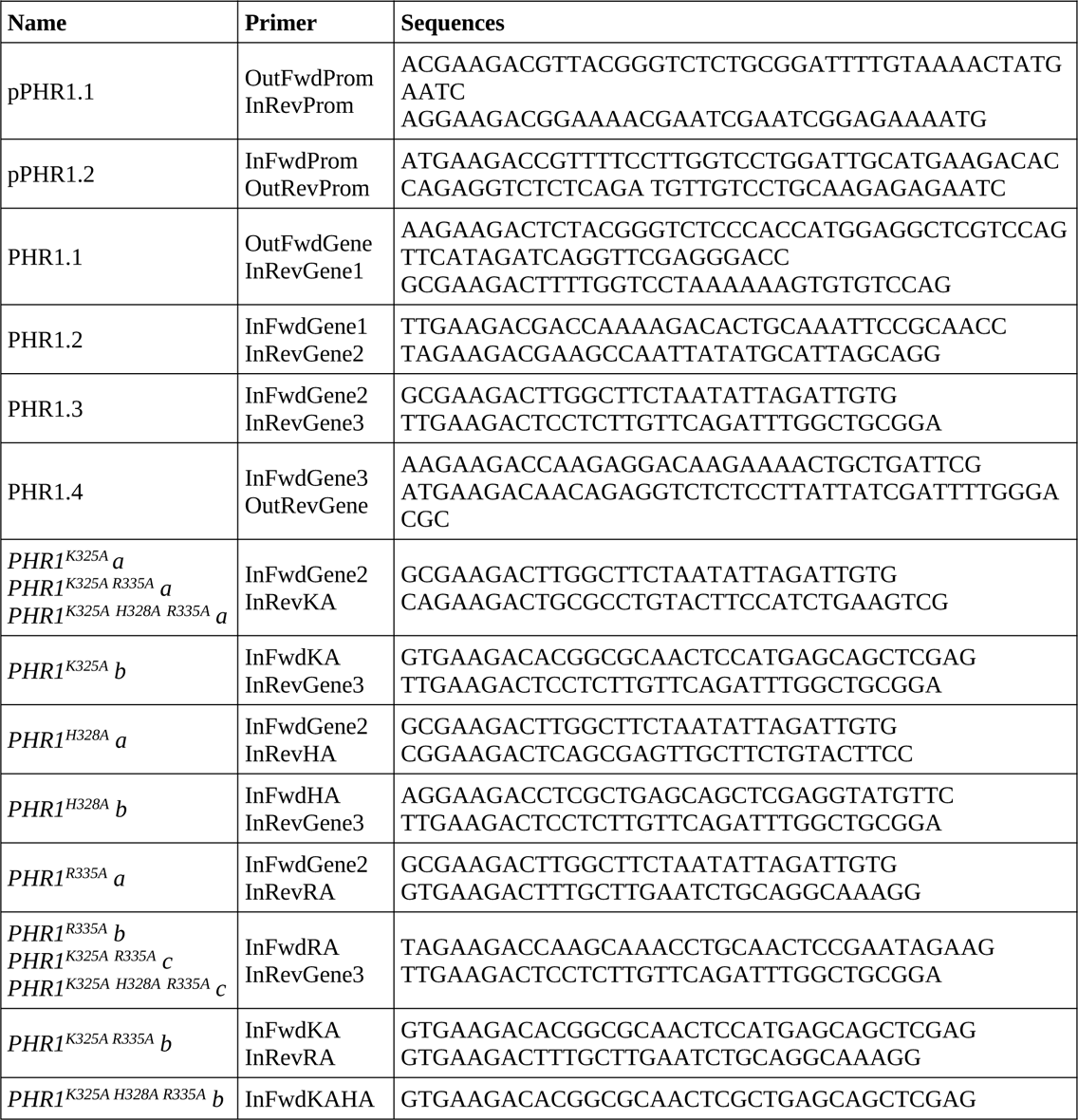
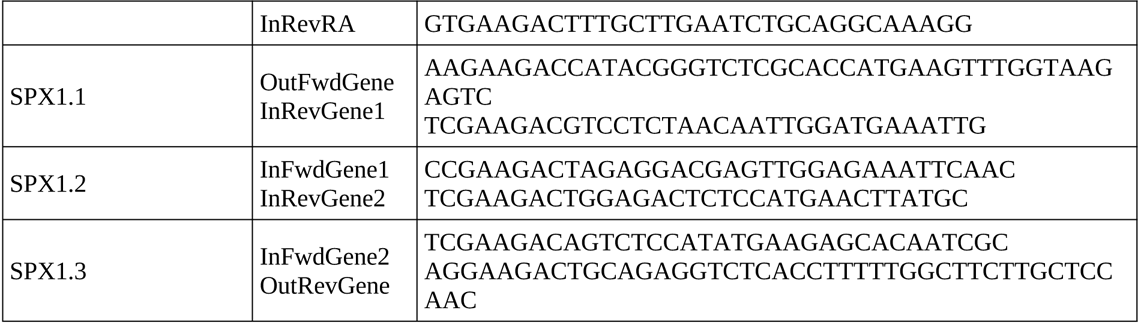
– Constructs and primers. **a, Golden Gate Level 0 constructs and primers.** The *PHR1* promoter and gene were amplified from *A. thaliana* gDNA, and *SPX1* was amplified from *A. thaliana* cDNA. Level 0 constructs were generated via StuI or SmaI cut-ligation into pUC Amp^70^.

**b, Level I, II & III constructs.** Level 1 and level 3 constructs were generated via BpiI cut-ligation, and level 2 constructs via BsaI cut-ligation^70^.

| Name | Assembly | Purpose |
| --- | --- | --- |
| LI BpiI pPHR1 | pPHR1.1 + pPHR1.2 | Cloning |
| LI BpiI PHR1 | PHR1.1 + PHR1.2 + PHR1.3 + PHR1.4 | Cloning |
| *LI BpiI PHR1<sup>K</sup>* | PHR1.1 + PHR1.2 + PHR1<sup>K325A</sup> a + PHR1<sup>K325A</sup> b + PHR1.4 | Cloning |
| *LI BpiI PHR1<sup>H</sup>* | PHR1.1 + PHR1.2 + PHR1<sup>H328A</sup> a + PHR1<sup>H328A</sup> b + PHR1.4 | Cloning |
| *LI BpiI PHR1<sup>R</sup>* | PHR1.1 + PHR1.2 + PHR1<sup>R335A</sup> a + PHR1<sup>R335A</sup> b + PHR1.4 | Cloning |
| *LI BpiI PHR1<sup>KR</sup>* | PHR1.1 + PHR1.2 + PHR1<sup>K325A R335A</sup> a + PHR1<sup>K325A R335A</sup> b + PHR1<sup>K325A R335A</sup> c + PHR1.4 | Cloning |
| *LI BpiI PHR1<sup>KHR</sup>* | PHR1.1 + PHR1.2 + PHR1<sup>K325A H328A R335A</sup> a + PHR1<sup>K325A H328A R335A</sup> b + PHR1<sup>K325A H328A R335A</sup> c + PHR1.4 | Cloning |
| LI SPX1 | SPX1.1 + SPX1.2 + SPX1.3 | Cloning |
| *LII R5-6 pPHR1:FLAG-PHR1* | LI A-B pPHR1 + LI FLAG B-C + LI C-D PHR1 + LI dy D-E + LI E-F nos-T + LI dy F-G | Cloning |
| *LII R5-6 pPHR1:FLAG-PHR1<sup>K</sup>* | LI A-B pPHR1 + LI FLAG B-C + LI C-D PHR1<sup>K</sup> + LI dy D-E + LI E-F nos-T + LI dy F-G | Cloning |
| *LII R5-6 pPHR1:FLAG-PHR1<sup>H</sup>* | LI A-B pPHR1 + LI FLAG B-C + LI C-D PHR1<sup>H</sup> + LI dy D-E + LI E-F nos-T + LI dy F-G | Cloning |
| *LII R5-6 pPHR1:FLAG-PHR1<sup>R</sup>* | LI A-B pPHR1 + LI FLAG B-C + LI C-D PHR1<sup>R</sup> + LI dy D-E + LI E-F nos-T + LI dy F-G | Cloning |
| *LII R5-6 pPHR1:FLAG-PHR1<sup>KR</sup>* | LI A-B pPHR1 + LI FLAG B-C + LI C-D PHR1<sup>KR</sup> + LI dy D-E + LI E-F nos-T + LI dy F-G | Cloning |
| *LII R5-6 pPHR1:FLAG-PHR1<sup>KHR</sup>* | LI A-B pPHR1 + LI FLAG B-C + LI C-D PHR1<sup>KHR</sup> + LI dy D-E + LI E-F nos-T + LI dy F-G | Cloning |
| LII F1-2 p35S:mCherry | LI A-B p35S + LI dy B-C + LI C-D mCherry + LI dy D-E + LI E-F 35S-T + LI dy F-G | Cloning / *N. benthamiana* transformation |
| LII F1-2 p35S:mCherry-SPX1 | LI A-B p35S + LI mCherry B-C + LI C-D SPX1 + LI dy D-E + LI E-F nos-T + LI dy F-G | *N. benthamiana* transformation |
| LII F1-2 p35S:GFP-PHR1 | LI A-B p35S + LI GFP B-C + LI C-D PHR1 + LI dy D-E + LI E-F nos-T + LI dy F-G | *N. benthamiana* transformation |
| LII F1-2 p35S:GFP-PHR1<sup>KHR</sup> | LI A-B p35S + LI GFP B-C + LI C-D PHR1<sup>KHR</sup> + LI dy D-E + LI E-F nos-T + LI dy F-G | *N. benthamiana* transformation |
| LIIIβ fin p35S:mCherry pPHR1:FLAG-PHR1 | LII p35S:mCherry 1-2 + LII ins 2-3 + LII dy 3-4 + LII ins 4-5 + LII F LII pPHR1:FLAG-PHR1 5-6 | *A. thaliana* transformation |
| LIIIβ fin p35S:mCherry pPHR1:FLAG-PHR1<sup>K</sup> | LII p35S:mCherry 1-2 + LII ins 2-3 + LII dy 3-4 + LII ins 4-5 + LII F LII pPHR1:FLAG-PHR1<sup>K</sup> 5-6 | *A. thaliana* transformation |
| LIIIβ fin p35S:mCherry pPHR1:FLAG-PHR1<sup>H</sup> | LII p35S:mCherry 1-2 + LII ins 2-3 + LII dy 3-4 + LII ins 4-5 + LII F LII pPHR1:FLAG-PHR1<sup>H</sup> 5-6 | *A. thaliana* transformation |
| LIIIβ fin p35S:mCherry pPHR1:FLAG-PHR1<sup>R</sup> | LII p35S:mCherry 1-2 + LII ins 2-3 + LII dy 3-4 + LII ins 4-5 + LII F LII pPHR1:FLAG-PHR1<sup>R</sup> 5-6 | *A. thaliana* transformation |
| LIIIβ fin p35S:mCherry pPHR1:FLAG-PHR1<sup>KR</sup> | LII p35S:mCherry 1-2 + LII ins 2-3 + LII dy 3-4 + LII ins 4-5 + LII F LII pPHR1:FLAG-PHR1<sup>KR</sup> 5-6 | *A. thaliana* transformation |
| LIIIβ fin p35S:mCherry pPHR1:FLAG-PHR1<sup>KHR</sup> | LII p35S:mCherry 1-2 + LII ins 2-3 + LII dy 3-4 + LII ins 4-5 + LII F LII pPHR1:FLAG-PHR1<sup>KHR</sup> 5-6 | *A. thaliana* transformation |

**c, Plasmids for recombinant protein expression in *E. coli*.** Plasmids for recombinant protein expression in *E. coli* have been generated via Gibson cloning^68^. Mutations targeting AtPHR1 K325, H328 and R335 were introduced by site-directed mutagenesis PCR^69^.

| Vector | Construct | Purpose |
| --- | --- | --- |
| pMH_HT | *AtPHR1*<sup>222-358</sup> | OmniSEC, GCI, EMSA |
| pMH_HT | *AtPHR1*<sup>222-358</sup> Olig1 | OmniSEC, GCI, EMSA |
| pMH_HT | *AtPHR1*<sup>222-358</sup> Olig2 | OmniSEC, GCI, EMSA |
| pMH_HT | *AtPHR1*<sup>280 – 360</sup> | OmniSEC |
| pMH_HT | *AtPHR1*<sup>280 – 360</sup> Olig1 | OmniSEC |
| pMH_HT | *AtPHR1*<sup>280 – 360</sup> Olig2 | OmniSEC |
| pMH_HT | *AtPHR1*<sup>280-360 K325A, H328A, R335A</sup> | OmniSEC |
| pMH_HSgb1T | *OsPHR2*<sup>1 – 426</sup> | OmniSEC, GCI, ITC |
| pMH_HSgb1T | *OsPHR2*<sup>1 – 426 KHR</sup> | OmniSEC, GCI, ITC |
| pMH_Hssumo | *OsSPX4*<sup>1-321</sup> | ITC |
| pMH_Hssumo | *AtPHR1*<sup>280 – 360</sup> | Crystallisation |

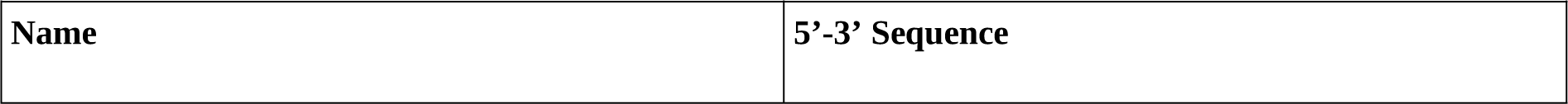
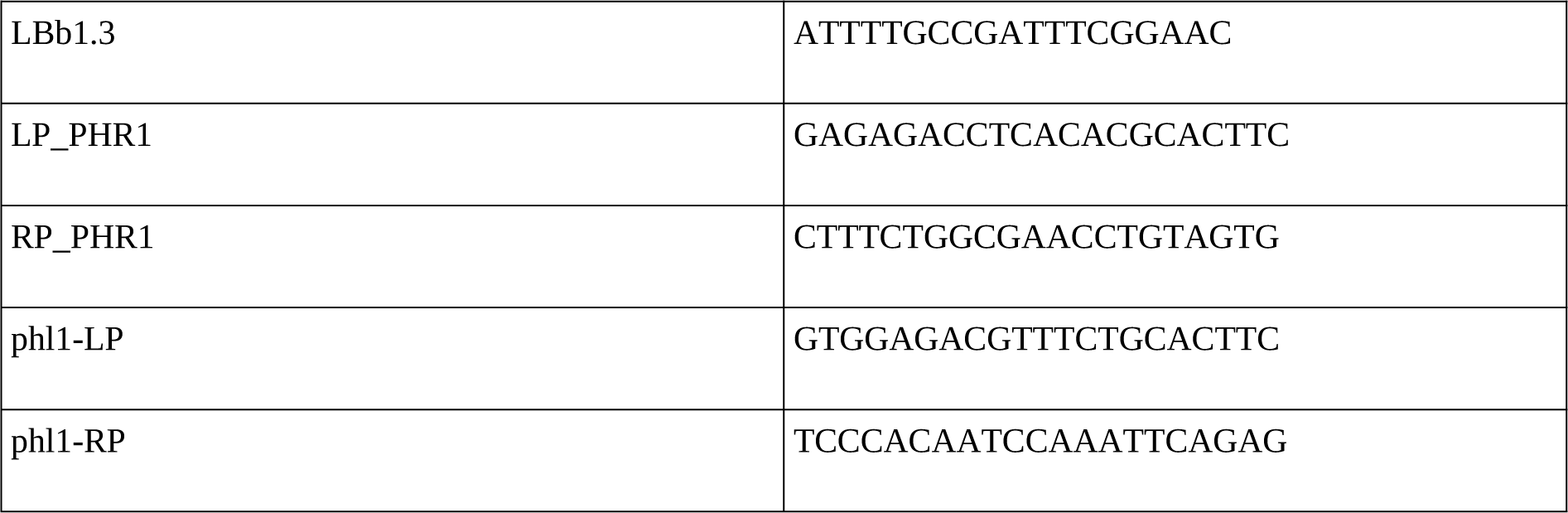
d, Characterisation of T-DNA mutants.

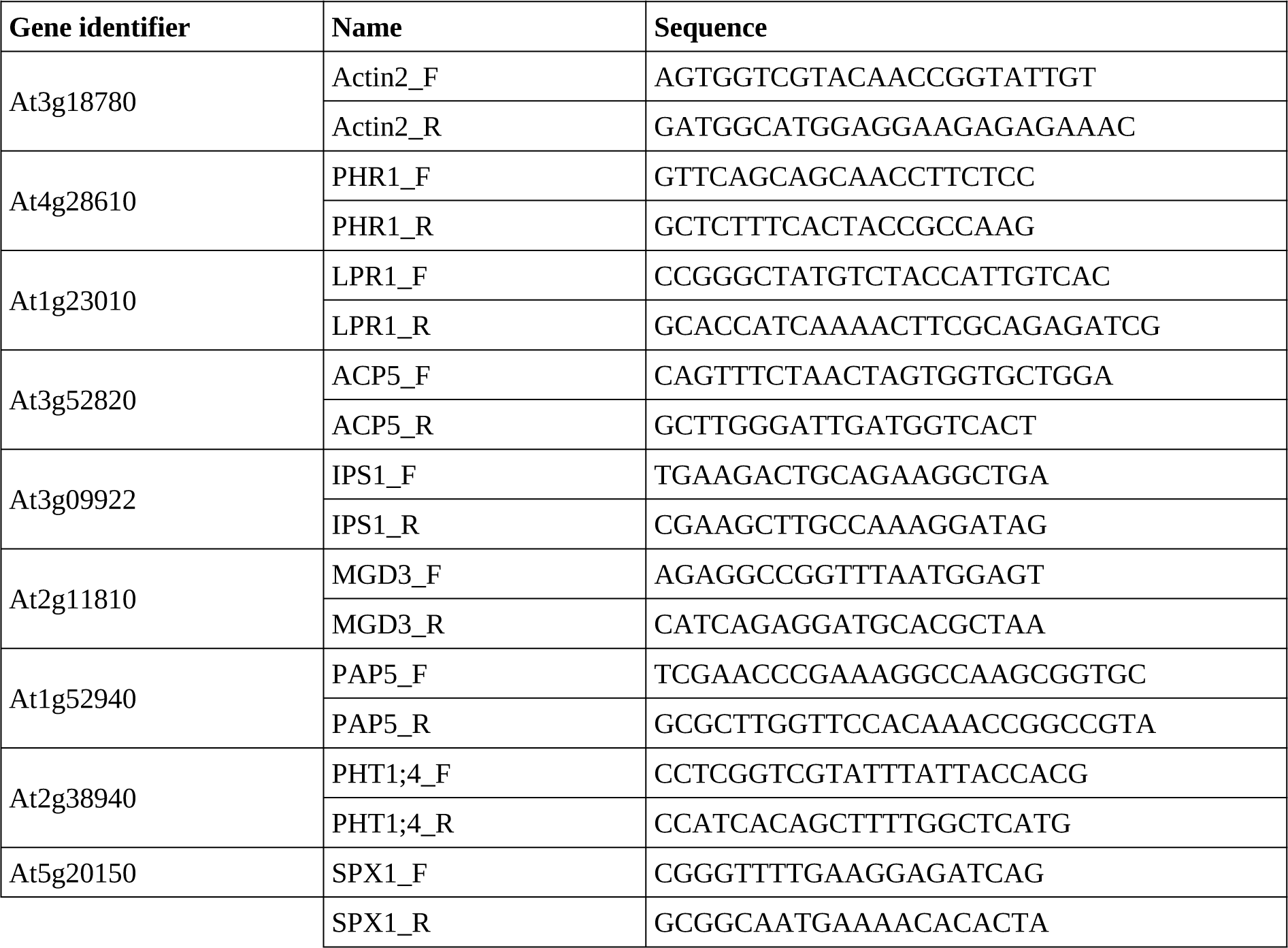
e, Gene expression analysis.

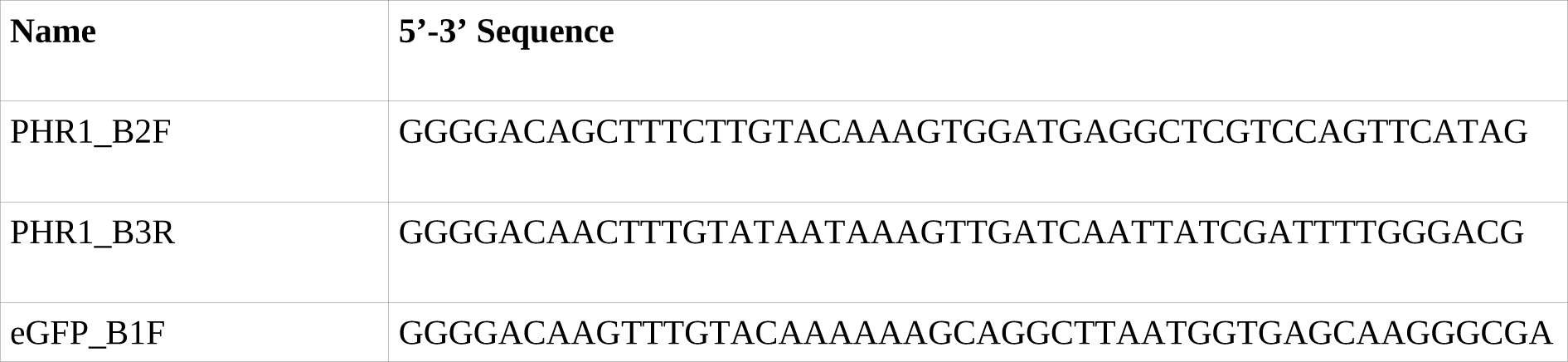
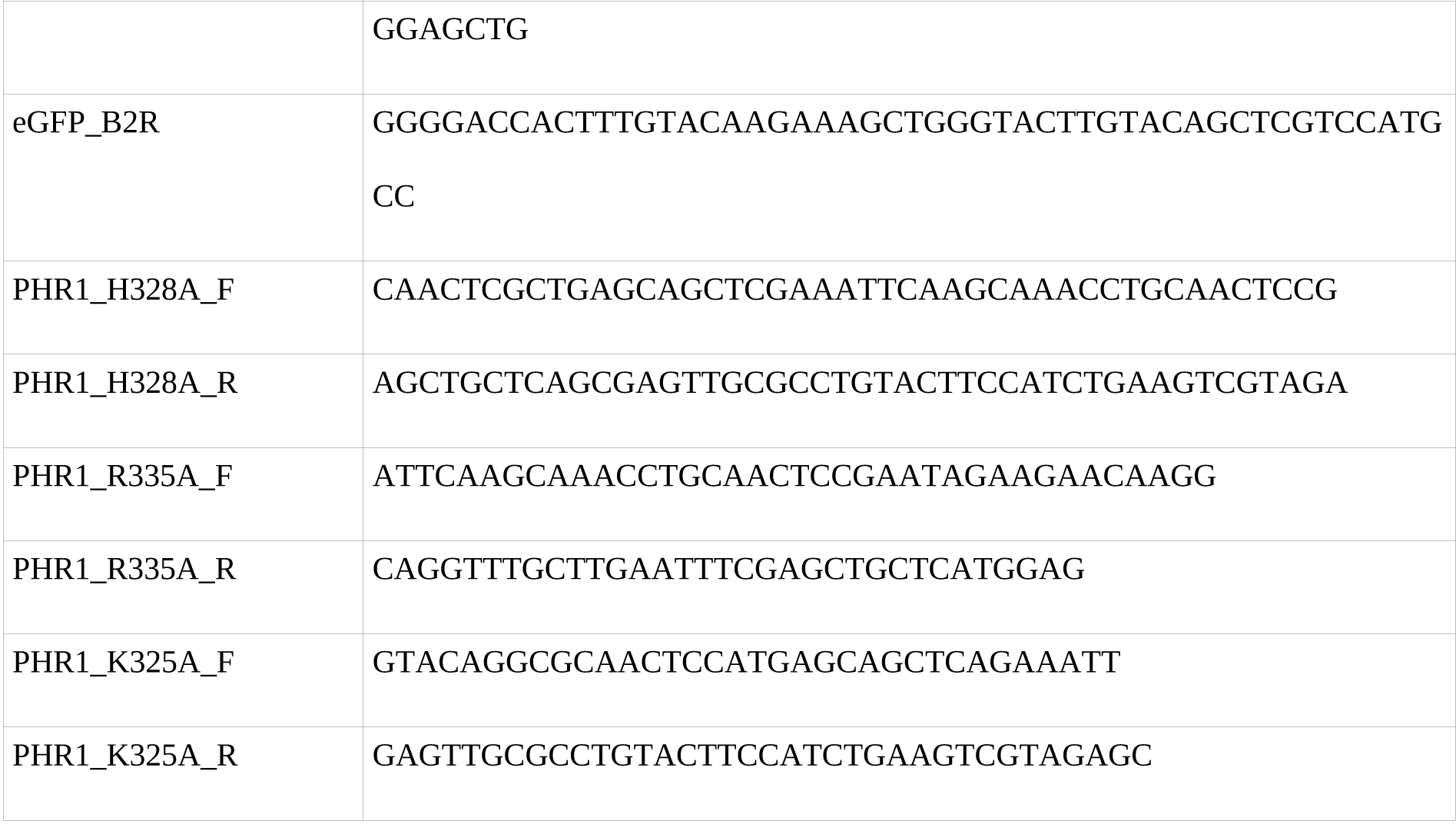
f, Primers used for cloning PHR1 into the pH7m34GW vector.

**Tabel 4.**
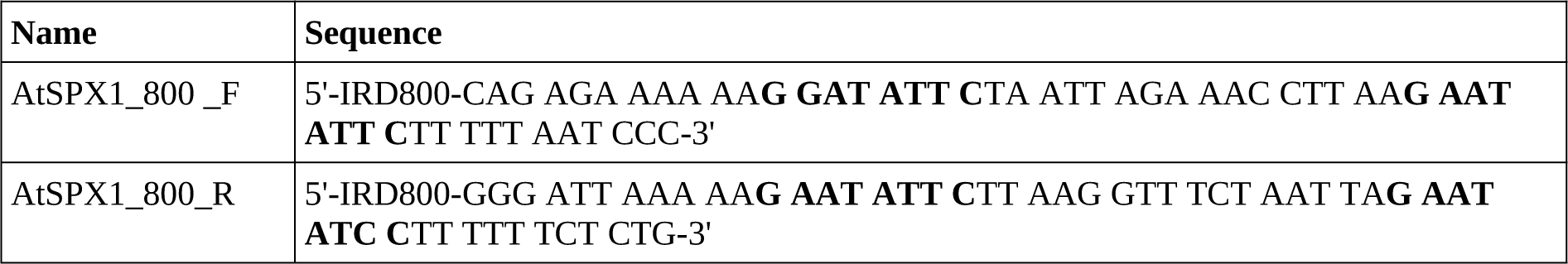
**Modified DNA oligos.** The P1BS (GNATATNC) is shown in bold. a, IRdye end-labelled oligos for EMSA.

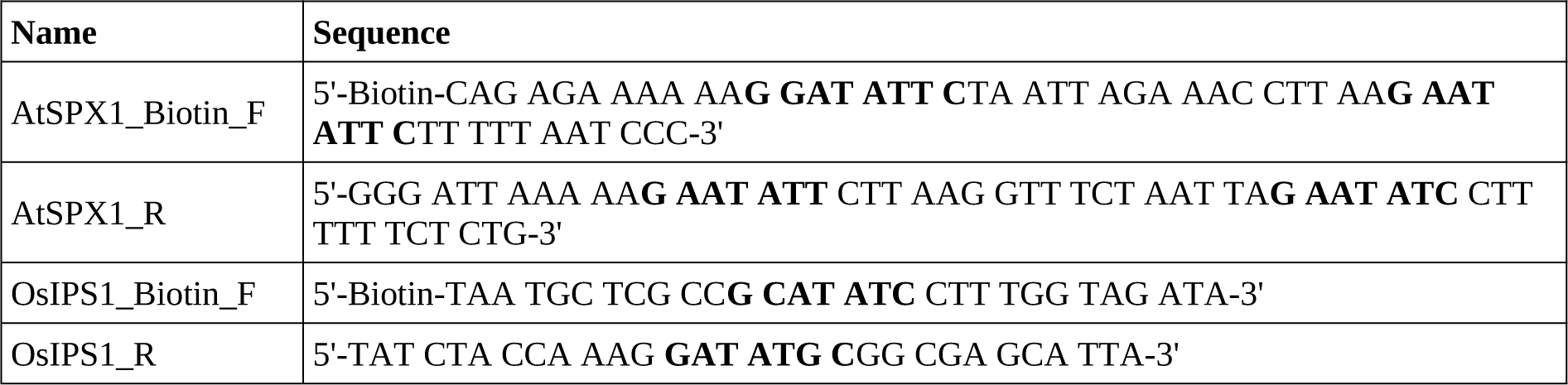
b, Biotinylated oligos for GCI.

**Supplementary Table 5.** – Crystallographic data collection and refinement statistics.

| PDB-ID | AtPHR1<sup>280-360</sup> form1<br>6TO5 | AtPHR1<sup>280-360</sup> form2<br>6TO9 | AtPHR1<sup>280-360</sup> form3<br>6TOC |
| --- | --- | --- | --- |
| **Data collection** |  |  |  |
| Space group | P 6<sub>1</sub> 2 2 | P 3<sub>2</sub> 2 1 | P 4<sub>2</sub> |
| Cell dimensions |  |  |  |
| *a*, *b*, *c* (Å) | 70.4, 70.4, 148.88 | 70.05, 70.05, 80.17 | 31.52, 31.52, 81.60 |
| α, β, γ (°) | 90, 90, 120 | 90, 90, 120 | 90, 90, 90 |
| Resolution (Å) | 47.18 – 2.38 (2.52 – 2.38) | 48.37 – 2.45 (2.59 – 2.44) | 31.52 – 1.85 (1.97 – 1.85) |
| *R*<sub>meas</sub><sup>#</sup> | 0.189 (2.21) | 0.164 (2.77) | 0.073 (2.76) |
| CC(1/2)<sup>#</sup> | 0.99 (0.69) | 0.99 (0.51) | 1.0 (0.40) |
| *I*/σ *I*<sup>#</sup> | 14.85 (1.48) | 15.86 (1.07) | 21.41 (0.98) |
| Completeness (%)<sup>#</sup> | 99.8 (98.9) | 99.3 (96.0) | 99.9 (99.3) |
| Redundancy<sup>#</sup> | 20.8 (20.2) | 19.0 (18.4) | 13.5 (13.4) |
| Wilson B-factor<sup>#</sup> | 56.7 | 67.8 | 47.1 |
| **Refinement** |  |  |  |
| Resolution (Å) | 41.18 – 2.38 | 48.37 – 2.45 | 31.52 – 1.85 |
| No. reflections | 16,560 | 8,702 | 6,433 |
| *R*<sub>work</sub>/*R*<sub>free</sub><sup>\$</sup> | 0.22 (0.23) | 0.23 (0.26) | 0.21 (0.26) |
| No. atoms |  |  |  |
| protein | 940 | 952 | 753 |
| solvent | 18 | 4 | 27 |
| Res. B-factors<sup>\$</sup> | | | |
| protein | 64.6 | 79.1 | 41.3 |
| solvent | 57.4 | 59.3 | 40.7 |
| R.m.s deviations<sup>\$</sup> | | | |
| bond lengths (Å) | 0.89 | 0.98 | 1.61 |
| bond angles (°) | 0.0047 | 0.0049 | 0.012 |
| Ramachandran plot<sup>\$</sup>: | | | |
| most favored regions (%) | 99.08 | 99.07 | 98.8 |
| outliers (%) | 0 | 0 | 0 |
| MolProbity score<sup>\$</sup> | 1.31 | 1.19 | 1.24 |

<sup>#</sup>as defined in XDS<sup>52</sup>
<sup>+</sup>as defined phenix.refine<sup>71</sup> (form 1, form2) or Refmac5<sup>56</sup> (form 3, using twin laws h,k,l and -k, -h, -l with twin fractions of 0.5, apparent point group was P 4 2 2)
<sup>\$</sup>as defined in Molprobity<sup>72</sup>

